# Changes in benthic and pelagic production interact with warming to drive responses to climate change in a temperate coastal ecosystem

**DOI:** 10.1101/2022.06.20.496925

**Authors:** Asta Audzijonyte, Gustav Delius, Rick D. Stuart-Smith, Camilla Novaglio, Graham J. Edgar, Neville S. Barrett, Julia L. Blanchard

## Abstract

Changing sea temperatures and primary productivity are rapidly altering marine ecosystems, but with considerable uncertainty in our understanding of the relative importance of these drivers and how their interactions may affect fisheries yield through complex food webs. Such outcomes are more difficult to predict for shallow coastal ecosystems than those in pelagic and shelf habitats, because coastal food webs are fuelled by a combination of separate pelagic and benthic energy pathways. Using long-term, empirical field data, we developed a novel multispecies size spectrum model for shallow coastal reefs. We include size-structured benthic and pelagic resources and trophic structures, allowing us to explore potential climate change scenarios that involve varying combinations of warming with changes in benthic and pelagic resources. Our model predicts that changes in resource levels will have much stronger impacts on fish biomass and yields than changes driven by physiological responses to temperature. Under increased plankton abundance, species in all trophic groups were predicted to increase in biomass, average size and yields. By contrast, changes in benthic resource produced variable responses across coastal trophic groups. Increased benthic resource led to increasing benthivorous and piscivorous fish biomass, yields and mean body sizes, but decreases in herbivores and planktivores. When resource changes were combined with warming seas, physiological responses generally decreased species’ biomass and yields. Our results suggest that the source, size and abundance of primary and secondary producers are critical to understanding impacts of warming seas on coastal fish communities. Understanding changes in benthic production and its implications for coastal fisheries requires urgent attention. Our modified size spectrum model provides a framework for further study of benthic and pelagic energy pathways that can be easily adapted to other ecosystems.

## Introduction

Climate change is causing a rapid and poorly understood reorganisation of natural ecosystems worldwide (Doney et al. 2012, Poloczanska et al. 2016), with implications for human wellbeing and conservation goals (Scheffers et al. 2016, Thiault et al. 2019). Shifts in species distributions, altered primary production, and changing physiological rates have been documented in terrestrial and aquatic ecosystems and are re-shaping food webs (Brose et al. 2012, Gibert and DeLong 2014, Kortsch et al. 2015). Understanding, predicting and mitigating consequences of these changes on food production and biodiversity conservation represent important research priorities (Blanchard et al. 2012), but predictions remain extremely difficult due to the complexity of interactions (Payne et al. 2016). One of the main challenges in attempting to predict responses of the valuable fishes in marine ecosystems is that changing ocean climates can impact fish populations through changes in primary productivity, the size distributions of their food resources, fish community composition and size structure, and the influences of temperature on growth and other metabolic processes (Boyd et al. 2015, Woodworth-Jefcoats et al. 2019). To understand how these changes will affect food webs, we need tools that can resolve physiological processes, body size structures and species interactions dynamically, in a robust and computationally efficient way.

Physiologically structured food web models are particularly useful for this purpose because they incorporate key organismal processes that likely respond to temperature, resolve size structure interactions between organisms, and allow species abundance, interactions, growth and yields to emerge dynamically from the assumptions and knowledge about species life-history, density dependence and spatial overlap (Andersen 2019). Such modelling tools have already been applied to explore climate change impacts on marine ecosystems at regional and global level, and have predicted declining fisheries yields, decreasing energy transfer efficiency, decreasing growth rates, and smaller fish body sizes with warming (Blanchard et al. 2012, Lefort et al. 2015, Bryndum-Buchholz et al. 2019, Lotze et al. 2019, Woodworth-Jefcoats et al. 2019). Regardless of these advances, climate and productivity change impacts remain highly uncertain, especially for coastal regions, where the relative contributions of pelagic versus benthic primary production are largely unknown (Brown et al. 2010, Everett et al. 2020). An additional consideration is that fish populations are affected both by overall productivity shifts and by changing food resource size composition (Woodworth-Jefcoats et al. 2013).

In most food web models, exploring ecological impacts of climate change under future climate scenarios, projected productivity and temperature changes are applied together, typically by forcing these models with temperature and production time series generated by Earth System Models (e.g. Coupled Model Intercomparison Project), and concentrating on long-term projections, summarising results across the entire community (e.g. (Bryndum-Buchholz et al. 2019, Tittensor et al. 2021)). While needed for large scale predictions, this approach makes it hard to understand the relative importance and uncertainties associated with production, temperature, size structure and their interactions. An alternative approach is to examine productivity, resource size structure, and temperature treatments separately and combined to better understand the mechanisms and interactions driving changes (Heneghan et al. 2021b).

A major challenge for predictions in coastal ecosystems is that global Earth Systems Models do not yet resolve benthic primary production pathways that are likely to play important roles for coastal communities. Even though global ecological models that capture ecological interactions between simplified benthic and pelagic pathways do exist for climate projections (Blanchard et al. 2012, Petrik et al. 2020), the energy pathways that fuel their production are not well resolved. Yet, shallow water marine ecosystems are undergoing some of the most rapid and accelerating changes due to human impacts (Lotze et al. 2006, Halpern et al. 2008), provide livelihood for over one billion people, and harbour most of marine biodiversity given that coral reefs are estimated to include ~25% of all marine species on their own (Fisher et al. 2015).

Shallow water systems differ from open ocean in several important ways, and models developed for open ocean systems may be limited in their capacity to make predictions for shallow ecosystems. First, light penetrates to the substrate in shallow coastal ecosystems, supporting an additional primary production pathway through benthic producers. The benthic production pathway encompasses production by microphytobenthos, tiny turf algae, and macroalgae up to giant kelp or a diverse array of corals, fuelling a multitude of benthic invertebrate consumers and predators (Duffill Telsnig et al. 2019, Chen et al. 2021). The relative contributions of benthic and pelagic pathways in coastal ecosystems remains debated (Docmac et al. 2017) and likely varies in time and space, yet it seems reasonable to expect that climate change will impact these two pathways in different ways. Second, higher structural complexity in coastal ecosystems represents an important difference to open ocean and shelf ecosystems, influencing the transferability of model-based predictions. The habitat and resource complexity provided by corals and kelps growing on rocky and coral reefs provides important refugia from predation (Rogers et al. 2014) and supports higher functional diversity of reef trophic groups able to capitalise on the diverse range of resources (Stuart-Smith et al. 2013). Unlike in open ocean or shelf systems (e.g. (Blanchard et al. 2014, van Denderen et al. 2018, Andersen 2019), many fishes on shallow reefs feed on low trophic level resources, rather than on each other, which suggests that changes in resource abundance or size structure might disproportionately impact the fish community (e.g. (Blanchard et al. 2010)).

To better understand climate change responses in coastal systems we need to extend physiologically structured food web models to represent the specific features of coastal ecosystems. We should also explore the impacts of temperature, resource abundance, and resource size structure in a systematic fashion to make predictions under a range of plausible scenarios. For this purpose we here outline a model parameterised and calibrated for a well-studied temperate rocky coastal reef system in a climate change hotspot (Fig. 1), where most of the trophic groups, including some large fish species, rely on benthic resources (Fig. 2). Using the model, we systematically explore how changes in abundance and size composition of pelagic and benthic resources interact with physiological responses of fishes to warming, and, as a consequence, how coastal fish species and community characteristics are affected. We ask two broad sets of questions: 1) Do changes in the pelagic and benthic production pathways have similar or opposing effects on biomass, average body size and yields of different trophic groups, and on the overall fish production? 2) How do species and community-level responses to plankton and benthos changes compare to their responses to warming? Do temperature-driven changes amplify or dampen changes due to food resource availability?

**Fig. 1.**
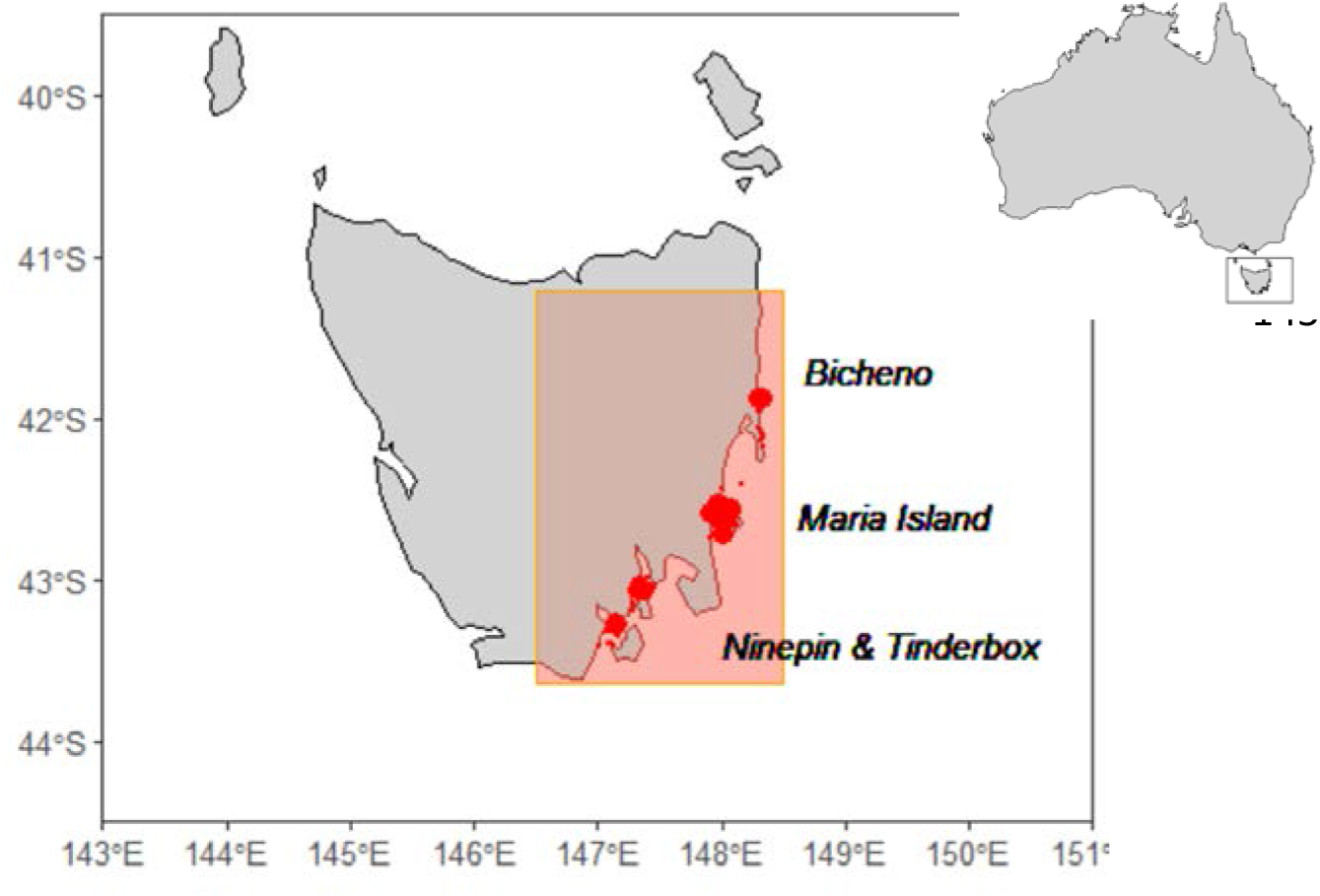
Area of the model domain in Tasmania (shaded in orange) and survey locations used to calibrate the model (circle area proportional to the number of surveys).

**Fig. 2.**
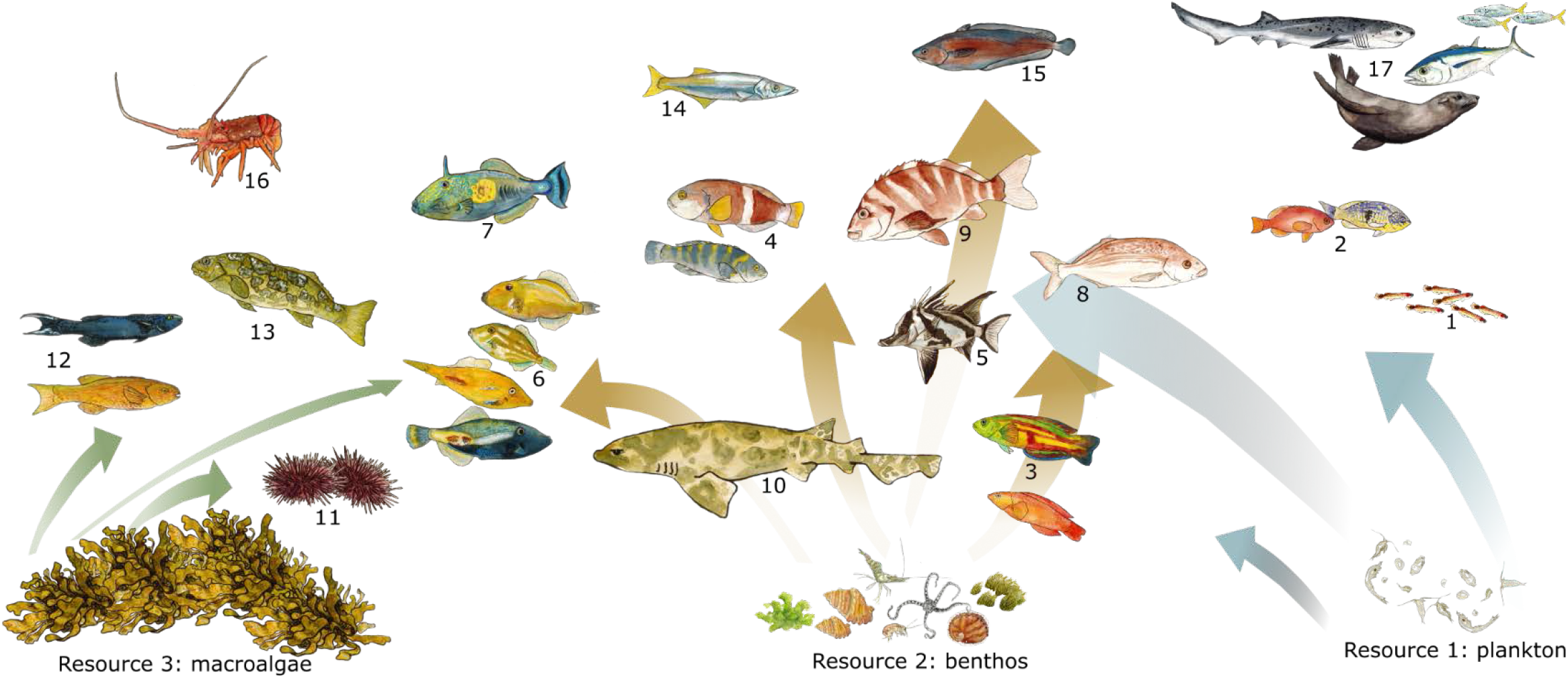
Schematic illustration of the model groups and three background resources used in this study. Detailed feeding interactions across species are not shown, coloured arrows indicate only the dominant feedings pathways for herbivores (left, from macroalgae), benthivores (middle, from benthos) and planktivores (right, from plankton). All species feed on plankton in earliest life stages (indicated with the large arrow). Predators (#14-17) are shown on the top of the figure, but for other groups position in the figure does not reflect their trophic level. The image was drawn by Amy Coghlan.

## Methods

### Model system and selection of taxa

To study the impacts of global warming on coastal fish communities, we focused on Tasmanian reefs within the SE Australian climate change hotspot, parameterising a multi-species size spectrum (MSSS) model for a well-studied temperate rocky reef community (Fig. 1). These communities have been monitored regularly for the last 30 years using standardised quantitative underwater visual surveys undertaken as part of the Australian Temperate Reef Collaboration (ATRC) monitoring program conducted since 1992 (Edgar and Barrett 2012), and Reef Life Survey (RLS) program since 2008 (Edgar and Stuart-Smith 2009, 2014), as described in an online methods manual at http://www.reeflifesurvey.com. In this study we focus on four long-term monitored locations, all positioned within a single biogeographical province. Data from the 1990s was used to parameterise the model, whereas data from 2000s and 2010s was used to compare empirical observations with alternative scenario outputs.

Selection of functional groups for inclusion in the model can have large impacts on the predicted ecosystem responses (Olivier and Planque 2017). To minimise the subjective choice of model groups we selected all species that satisfied the occurrence or biomass thresholds (see Supplement) in standard fish surveys (500m^2^ area, see (Edgar et al. 2016) for survey methods), and pooled species with similar life-history characteristics and diets. The final list comprised 14 fish and shark species and species groups (Table 1, Fig.2), which together accounted for over 90% of average observed biomass per survey in the 1990s. To account for the fact that abundance of mobile larger predators (marine mammals, tunas, mobile sharks, large predatory fishes, birds) is underestimated in underwater visual surveys, we also included a general large predator species to represent the unaccounted predation. Most of the smaller sized invertebrate species were represented through the benthic background resource, but two ecologically and economically important groups (urchins and rock lobsters) were modelled explicitly as dynamic size structured groups.

**Table 1.** List of model species and groups assigned into four main trophic categories, approximate maximum body size and modelled fishing mortality. Further details are given in the Supplement. * - the biomass of predators is not taken from surveys but assumed here to represent average predation from all large predators. Note, that fishing mortality in a continuous size-based model, such as the one used here, is not directly comparable to mortality in age-structured stock assessment models. Numbers next to common names refer to numbers in Fig. 2.

| Common name | Scientific name | $\sim W_{\infty}$<br>(kg) | F (year<sup>-1</sup>) |
| --- | --- | --- | --- |
| **Planktivores** |  |  |  |
| Hulafish (1) | *Trachinops caudimaculatus* | 0.04 | 0 |
| Barber Perch (2) | *Caesioperca rasor* | 0.62 | 0.05 |
| **Benthivores** |  |  |  |
| Senator wrasse (3) | *Pictilabrus laticlavus* | 0.40 | 0.05 |
| Purple & bluethroat wrasse (4) | *Notolabrus tetricus* & *Notolabrus fucicola* | 1.60 | 0.15 |
| Long-snouted boarfish (5) | *Pentaceropsis recurvirostris* | 2.20 | 0.15 |
| Leatherjackets (6) | *Acanthaluteres vittiger* & *Meuschenia australis* | 3.00 | 0.05 |
| Six-spine Leatherjacket (7) | *Meuschenia freycineti* | 4.30 | 0.15 |
| Bastard Trumpeter (8) | *Latridopsis forsteri* | 5.20 | 0.15 |
| Banded Morwong (9) | *Cheilodactylus spectabilis* | 13.50 | 0.15 |
| Draughtboard Shark (10) | *Cephaloscyllium laticeps* | 16.00 | 0.10 |
| **Herbivores** |  |  |  |
| Urchins (11) | *Heliocidaris*, *Centrostephanus* & *Goniocidaris* | 0.35 | 0.15 |
| Herring Cale (12) | *Olisthops cyanomelas* | 3.40 | 0.15 |
| Marblefish (13) | *Aplodactylus arctidens* | 4.40 | 0.15 |
| **Predators** |  |  |  |
| Long-fin Pike (14) | *Dinolestes lewini* | 1.00 | 0.15 |
| Red Cod (15) | *Pseudophycis palmata* | 1.50 | 0.15 |
| Rock lobsters (16) | *Jasus edwardsii* | 3.00 | 0.15 |
| Predators (general) (17) | Mammals, birds, sharks, tunas & other fish | 5.00 | 0.10 |

### General physiologically structured size spectrum modelling framework

To explore how temperature effects on fish physiology compare and interact with changes in the pelagic and benthic resource abundance and size composition, we use a multi-species size spectrum (MSSS) modelling framework (Andersen 2019) and its implementation in the R package *mizer* (Scott et al. 2014) (sizespectrum.org/mizer). The framework has been used in several recent studies (Blanchard et al. 2014, Spence et al. 2016, Reum et al. 2019a, Novaglio et al. 2021) and its theoretical basis is described in detail on sizespectrum.org, ‘mizer’ vignette and Andersen (2019). Here we only briefly mention the key assumptions, with more details provided in the Supplement.

The *mizer* application of MSSS model simulates a dynamic size-structured background resource and user-defined size structured groups (fish and invertebrates), which feed on the background and on each other. The groups can either represent a single species or groups of species and are referred to here as model groups. The dynamics of size structured groups is summarized by the McKendrik-von Foerster equation, where change in abundance at size through time depends on emergent somatic growth *g_i_*(*w*) [g year^-1^] and mortality μ_i_(*w*) [ year^-1^], which determines how many individuals enter and leave a size class respectively:

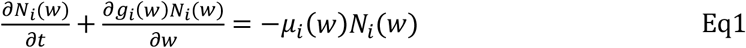

Growth depends on the availability of food, Holling type II feeding response, assimilation efficiency, maintenance costs, allocation to reproduction and growth efficiency (the latter is not typically used in other *mizer* applications). Mortality includes constant background mortality, predation, starvation, senescence and fishing mortality. The minimum, maximum and maturation sizes for each size-structured group are set by the user (Table 1, Tables S1, S2). The numbers in the first size class are determined by the continuous density-dependent recruitment (see below).

The temporal dynamics of the background resource spectrum (*N_R_*) is modelled through semi-chemostat dynamics (De Roos and Persson 2001) and depends on the emergent predation mortality (*μ*), resource regeneration rate (*r_0_*), and resource carrying capacity (*κ*) scaled by the size spectrum slope of the resource (*λ*):

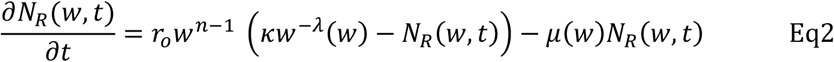

where *n* is the maximum food intake body scaling exponent of size structured groups (see (Hartvig et al. 2011) for further derivations of background spectra dynamics).

### Modification of the modelling framework for coastal ecosystems: benthic and pelagic production pathways

In this study we modified the standard MSSS model framework to suit coastal ecosystems by introducing three size-structured background spectra to represent pelagic, benthic (turf and invertebrates) and macroalgal (kelp, seaweed) resources. This distinction is important for shallow water communities, because: 1) a number of fish species are specialised to feed on either pelagic, benthic or macroalgal food and feeding on both benthic and pelagic food resources is size based (Brose et al. 2006), 2) in contrast to pelagic ecosystems, benthic primary and secondary production are likely to provide substantial energy inputs, independent from pelagic production (Jennings et al. 2001, Duffill Telsnig et al. 2019), 3) size spectrum slopes of pelagic and benthic producers, and their responses to climatic warming, are likely to differ, 4) many fish species have ontogenetic diet shifts, starting with pelagic (planktonic) prey and switching to benthic prey as they grow (Reum et al. 2019a); if these resources respond differently to warming seas, the implications for fishes and communities are likely to differ from those based on the assumption of a fixed resource for each species.

Ontogenetic diet shifts should ideally emerge dynamically in the model from assumptions about size preferences, habitat preference and relative food abundances, yet their reproduction in physiologically structured models can be challenging (Pethybridge et al. 2019, Reum et al. 2019a). In this study we aimed to reproduce emergent diet shifts by assuming different size ranges and size abundance slopes for the three resource spectra, and by specifying availability of each spectrum to each model species, implicitly representing habitat and food preference (pelagic, benthic, macroalgal). In this way each background resource has specific parameters for carrying capacity (*κ*), size spectrum slope (*λ*) and regeneration rate (*r_0_*), as well as minimum and maximum size of the spectrum (Table 2), whereas each species has resource-specific vectors indicating maximum availability of each resource (see below and Supplement). This means that emergent ontogenetic shifts could be reproduced by just three species-specific parameters, identifying preference for each of the three background resources. This modification of the *mizer* package, allowing multiple size-structured background spectra and species-specific preferences for different resources is now available as a *mizerMR* extension to the main *mizer* package (https://github.com/sizespectrum/mizerMR).

**Table 2.**
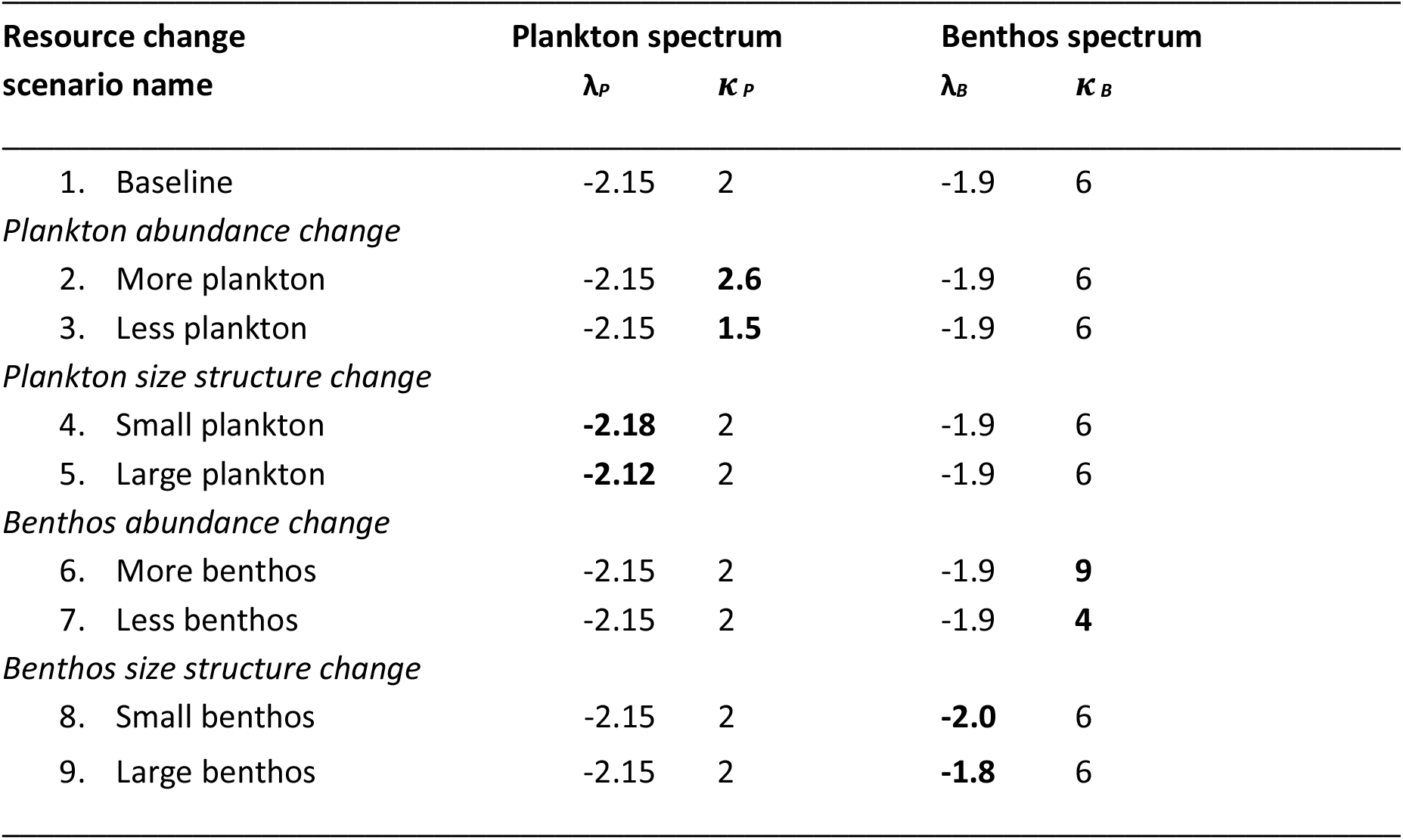
Scenarios of plankton and benthos resource changes. Each of these scenarios was combined with warming and fishing scenarios in a fully crossed ANOVA-style design.

### Climate change scenarios: changes in plankton and benthos abundance and size

To assess species and ecosystem responses to climate change scenarios, we explored nine alternative benthic and pelagic resource change scenarios (see below and Table 2) fully crossed with physiological temperature responses (i.e., temperature impacts on vital physiological rates) in an ANOVA-style design. For all scenarios low fishing mortality was assumed (Table 1). This gave a total of 18 scenarios, each run with 29 alternative parameter combinations to assess model output uncertainty (see below), resulting in 522 simulation outputs. Each scenario was initiated from baseline equilibrium conditions (no resource or temperature change, main parameter set), then new conditions imposed instantaneously and applied for 150 years of the model run. In nearly all cases simulations settled into a new stable or oscillating equilibrium within 20-40 years. Due to the large number of simulations, we did not focus on the transient dynamics, but compared final equilibrium conditions to the initial equilibrium state. To assess the impact of parameter uncertainty on predicted ecosystem responses in each of the 18 scenarios, we compared the differences between baseline and specific scenarios for each of the 29 parameter combinations respectively. This means that we were not focused on the total amount of uncertainty that combines both parameter and scenario uncertainty, but on whether a similar change in the modelled community occurred between the baseline and a specific model scenario, regardless of parameter combination.

#### Resource parameters in baseline and climate change scenarios

For many marine ecosystems, the slope of the plankton normalised abundance spectrum varies between −2 and −2.2 (Andersen 2019; Atkinson et al. 2020). For low productivity waters, such as SE Australia, slopes are often steeper and therefore for the baseline scenario we use the value of −2.15 (see Supplement for further details on background parameter values). Slopes and abundances of the benthos spectrum for the baseline scenario were estimated from empirical data along SE Australian coast (Supplement). The macroalgal spectrum was modelled here as size structured background resource for simplicity, even though herbivore feeding on macroalgae is unlikely to be primarily size based. To ensure high abundance of the macroalgal resource across various size groups, we assumed relatively flat slopes (−1.6) and high abundance (16g/m^2^). These values do not account for the large kelp stands in many rocky reef communities, but entire kelp is not typically consumed by the local herbivores.

The goal of our study was to explore consequences of changes in pelagic and benthic resource abundance and size structure. We were not aiming to replicate specific climate change scenarios because of large uncertainties in forecast changes for SE Australia and coastal ecosystems in general (Lotze et al. 2019, Tittensor et al. 2021). For example, it is generally expected that increasing temperature favours smaller body size and that higher temperatures will lead to steeper background resource size spectra (Reuman et al. 2008, Polovina and Woodworth 2012, Peter and Sommer 2013, Tittensor et al. 2021) (see Supplement). Yet empirical evidence shows that changes in temperature also affect nutrient availability and grazing pressure (Mazurkiewicz et al. 2020, Pomati et al. 2020), that benthic resource size spectra may be insensitive to temperature (Mazurkiewicz et al. 2020, Heather et al. 2021), and field data from the last 10 years shows increasing zooplankton abundance off Maria Island (Everett et al. 2020). We therefore explored a range of resource change scenarios, with plankton or benthos abundance (*κ*) and slopes (*λ*) increasing or decreasing independently (Table 2). We assumed that plankton or benthos *κ* values increase or decrease by ~30%, and slopes for plankton can change by 0.03 and for benthos by 0.1 (see Supplement).

#### Physiological response in climate change scenarios

To model temperature effects on physiology in a consistent manner with other similar physiologically structured models, we followed the metabolic theory of ecology approach (Brown et al. 2004) and assumed that temperature affects the rate of metabolism, search rate, maximum food intake rate, and background and senescence mortality (Pauly 1980; Englund *et al*. 2011; Rall *et al*. 2012). All rates were assumed to increase exponentially with temperature based on Arrhenius temperature correction factor

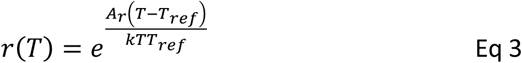

where *A_r_* is the activation energy [eV] for individual rate *r* here assumed to be 0.63, *T* is temperature [K], *T_ref_* is the reference temperature where temperature scaling equals 1 (here assumed to be 12°C or here 285.27 K), and *κ* is Boltzmann’s constant in eV K^-1^ ( 8.617 × 10^-5^ eV K^-1^). This representation of temperature is a simplification (see Discussion), and likely does not reflect inter-generational population physiological responses (Denderen et al. 2020, Wootton et al. 2022). Yet, it allows consistent comparison with other models exploring climate change effects (Blanchard et al. 2012, Bryndum-Buchholz et al. 2019, Heneghan et al. 2021a), results in a tractable number of simulations for ANOVA-type analyses, and, in a manageable way, addresses relative importance of internal, physiological versus external, community driven, responses to warming. As with productivity, changes in temperature were applied instantaneously to the baseline equilibrium conditions and the model run to a new equilibrium to assess species responses. We follow predictions of RCP8.5 scenario, forecasting temperature increase of ca 2.5°C by 2100 for SE Australia (Supplement), and temperature increase scenarios are run at 14.5°C temperature, compared to 12°C reference temperature. The selected parameter values lead to ca 20% increase in physiological rates.

### Species parameters and model parameterisation

Species parameters were selected based on either general size spectrum model assumptions (Hartvig et al. 2011, Andersen 2019), or estimated for this model (Supplement). For many rocky reef species maturation sizes and growth curve parameters are not known, hence we could not use standard growth based approaches to estimate intake and metabolism parameters (e.g. (Blanchard et al. 2014, Andersen 2019)), but instead estimated them from general inter-specific relationships (Charnov et al. 2013, Andersen 2019) or derived them from broad scale species correlations and estimates used in the Dynamic Energy Budget database (see Supplement, Fig. S1 and https://github.com/astaaudzi/SEAmodel).code for full details). To our knowledge, this is the first study providing alternative multi-species parameter derivations for MSSS models, and thus a novel test example for new model developments in data poor regions. Parameters defining intake rates were further explored in the uncertainty evaluation framework (below).

In addition to species specific life-history and physiological parameters, food web models are also highly sensitive to parameters defining species interactions. In MSSS models, the emergent consumption of a species or a background resource depends on its relative abundance at size, the consumer predation kernel (Table S1, Eq3) that sets limits on the size ranges that each species can eat, and the species interaction matrix (Table S1, Eq 14) that defines the maximum proportion of a prey species or resource biomass available for consumption at each time step. Predation kernel parameters were selected to reflect general expert knowledge and evidence from earlier studies (Blanchard et al. 2014, Soler et al. 2016)) (Table S4). For resource consumption, the availability of each background resource for each species was set through species and resource specific scalars, selected on the basis of general knowledge about functional groups (e.g. planktivores, herbivores, invertivores, predators (Stuart-Smith et al. 2013)), and aimed to achieve realistic emergent species diets (Table S5, Fig. S2). Briefly, for the 17 species, the interaction matrix can have up to 17×17 parameters, defining specific predation preferences for each species pair. In spatially implicit models, such as the one assumed here, the interaction matrix can be complex and used to model species physical overlap e.g. (Blanchard et al. 2014) or detailed diet preferences (Reum et al. 2019a). In this study, where species have full spatial overlap, the interaction matrix was reduced to just 5 parameters aimed to reflect general diet preferences and vulnerability to predation. Two parameters (0 and 0.7) indicate absence or presence of predation, and the remaining three parameters were used to adjust vulnerability to predation through anti-predatory behaviour, e.g. schooling in small bodied species, or morphological defences in urchins (Supplement). The interaction matrix parameters were further explored in uncertainty analyses.

One of the key parts of model parametrisation and calibration is finding parameters for reproduction and recruitment that enable species co-existence, reasonable biomass values, and expected resilience to exploitation (Jacobsen et al. 2017). To this end, we iteratively tuned two species-specific reproduction parameters - the maximum recruitment parameter *R_max_* and the reproductive efficiency parameter *ε* (Eq10, 11 in Table S1). The *R_max_* sets the upper recruitment limit and represents density dependent processes that are not dependent on the biomass, such as habitat availability or disease. The parameter *ε* determines the proportion of total reproduction energy converted into egg biomass and is used to account for reproduction inefficiencies, costs and early egg mortality (Andersen 2019). In this way, *ε* allows a linear relationship between stock biomass and recruitment, whereas *R_max_* adds the non-linear density dependence on recruitment. The two parameters were tuned to satisfy two following conditions. First, *R_max_* was adjusted to ensure relative model biomasses for each species at equilibrium conditions were within 20% of the observed relative biomasses in visual surveys from the 1990s (across sites and years, Table S3); this also ensured species co-existence. Second, *ε* was adjusted so that individual species’ vulnerability to fishing and fishing mortality at maximum yield (F_msy_) was within the expected range given species body size and life-history traits (Andersen 2019) (Supplement and https://github.com/astaaudzi/SEAmodel).

### Parameter uncertainty evaluation

The parameter selection procedure above, leads to one set of parameters that generates realistic model behaviour. Most multi-species food web and complex models with large numbers of parameter apply a similar procedure (e.g. (Smith et al. 2015, Jacobsen et al. 2017, Bossier et al. 2018, Ayllón et al. 2021)), with more rigorous parameter uncertainty evaluation limited to specific cases (e.g. (Thorpe et al. 2015, Spence et al. 2016)) and mostly focusing on species recruitment parameters (*R_max_*) evaluated against time series of catches. In this study we conducted uncertainty evaluation of 37 parameters, focusing on recruitment and species interaction parameters that were found to be influential and highly uncertain during the model exploration: species-specific *R_max,i_* and search rates (*γ_i_*) and three species interaction matrix parameter defining vulnerability to predation (Supplement).

To apply a relatively straightforward uncertainty evaluation procedure that could be replicated in other MSSS models, we used a rejection algorithm, similar to the Approximate Bayesian Computation approach (Beaumont 2010). For this we ran simulations with many parameter combinations and rejected simulations that did not satisfy specific criteria on emergent community properties, given the knowledge about the system. More specifically, for the 37 explored parameters, we sampled 1.8 ×10^6^ combinations, multiplying the parameter vector by a random uniform vector ranging from 0.5 to 2 (i.e. each parameter value was allowed to vary independently two-fold from the initial calibrated value). To explore the space around the initial values more closely, an additional 0.4 ×10^6^ parameter combinations were sampled using a multiplier of 0.8 to 1.2 (values were allowed to vary by 20%). With each of the resulting 2.2 ×10^6^ parameter combinations we ran the model for 150 years using a timestep of 0.5 years (instead of 0.2 years) to speed up the calculations and starting each run using equilibrium model state from the initial ‘tuned’ set of parameters. Each run was rejected if, at the end of the 150-year simulation, biomasses of at least one species were 4 times smaller or larger than the average biomasses observed in underwater surveys. This criterion alone reduced the parameter space from 2.2 ×10^6^ to 287 ×10^3^ combinations. The remaining parameter values were used in the second step, where for each parameter combination we ran the simulation five times, saving additional model outputs to explore emergent ecosystem responses and imposing additional mortality on key model species (set as fishing mortality). These “extra mortality” runs included: 1) no extra mortality, 2) extra mortality on urchins, 3) extra mortality on “predator”, 4) extra mortality on lobsters, 5) extra mortality on the planktivore *Trachinops caudimaculatus.* These runs were assessed using seven criteria, describing general characteristics of an unfished system (feeding level, diets) or the system’s response to imposing extra mortality on key species (Supplement). Once all the criteria were applied, we were left with a set of 28 parameter combinations, which together with the initial set of parameters gave the final set of 29 parameter combinations. These were used to explore uncertainty around alternative scenarios. We emphasise that our protocol does not adequately explore full parameter space and cannot be treated as probabilistic statistical uncertainty evaluation; hence we do not use confidence intervals or posterior probability terms. However, it provides some understanding of variation in model outputs and system responses to perturbations under different plausible parameter values.

### Comparing alternative scenario outputs

Baseline model dynamics and emergent properties were compared to available data and knowledge on species biomasses, growth, and emergent diets (Supplement). For each scenario we calculated four characteristics for all model groups – biomasses, yields under constant fishing mortality (Table 1) and mean body weight for all individuals above 2 cm in length (same minimum size cut-off as in visual underwater surveys). Changes in species biomasses and mean body size were compared with field observations, where for biomasses we used mean biomass per survey from 2013-2018, and for body sizes we used estimated changes in mean body size from Bayesian mixed-effect modelling analyses (Audzijonyte et al. 2020). All response variables were assessed at equilibrium conditions (100 years after applying new resource and temperature parameters). Because some scenarios settled into oscillating equilibrium with ca 10y periodicity, statistics were calculated as averages of the last 30 years.

As individual species within each of the four trophic groups showed similar responses to projected resource changes, we assessed more general trophic group level responses using mixed-effect ANOVA analyses. Separate analyses were conducted for plankton and benthos slope (*λ*) and abundance (*κ*) scenarios, making four sets of analyses in total. For each analysis, variation between species was treated as a random effect, whereas resource change (*Re*) (−1, 0, 1), warming (*W*) (0, 1) and trophic group (*T*) were fixed effects. We explored different levels of interactions, starting from the full model with a three-way interaction, then reducing the complexity and assessing alternative models using Akaike Information Criterion (AIC) (Table S7). For nearly all analyses the best selected model included all two-way interactions *response ~ Re*\* *W* +*W*\**T* + *Re*\**T* + (1|species). Statistical analyses were conducted using R version 4.0.0 (R Core Team, 2020), the ‘effects’ (v4.1-4; Fox et al., 2019) and the ‘emmeans’ (v1.4.7; Lenth et al., 2020).

## Results

### Model parameterisation in the baseline scenario

Before applying climate change scenarios, we explored emergent model properties in the baseline scenario with fixed resource and temperature conditions, to check that model parameters reproduced reasonable system behaviour. Across the 29 accepted parameter combinations we observed a wide range of equilibrium biomasses, indicating that these parameters capture a range of alternative system states (Fig.S6); the modelled equilibrium biomasses were close to the observed interannual variability in species biomasses for the 1990s and 2000s. The emergent resilience to fishing, assessed as fishing mortality at maximum yield in equilibrium conditions, was within the expected range, based on species life-history characteristics (Table S6). Emergent diets in model groups reproduced expected ontogenetic shifts, where all species were initially feeding on plankton, with benthivores and herbivores switching to their respective resources, and predators moving from plankton to some benthos and to fish (Fig. S2). Some benthivore species had small fish in their adult diets, which was consistent with empirical observations (Soler et al. 2016). Emergent relative abundance of benthic resource was close to the available data on average abundance and biomass of benthic invertebrates across the east coast of Tasmania (Fig. S5).

### Species and trophic group responses to changes in resources abundance and temperature: biomasses and yields

When new resource levels or physiological temperature impacts were applied to the equilibrium biomasses of the baseline scenario, most species settled into new stable or oscillating equilibrium within 20-50 years. This suggested that species parameters (mostly the combination of *R_max_* and *ε*) used in the simulations could replicate dynamic species responses and density dependence adjustments to the community. Only three model groups were highly sensitive to the explored changes in resource with predicted biomass declining to extremely low levels (extinction) under some parameter combinations. These were the planktivorous serranid *Caesioperca rasor* and urchins for scenarios where plankton resource carrying capacity (*κ*) decreased or slope (*λ*) increased, and the wrasse *Pictilabrus laticlavius*, for scenarios with benthic resource changes (Fig. 3, Fig. S7). Therefore, for these species, uncertainty ranges in biomass and yield responses were very broad. The impact of increasing resource carrying capacity or steeper resource slopes had similar effects on the model group biomasses. This is because steepening resource slopes decreased resources abundance at largest sizes (> 1g) that were offset by complementary increases at smaller sizes (< 1g). Below we only discuss results from changes in resource abundance, whereas impacts of changing resource slope are shown in the Supplement (Fig. S7).

**Fig. 3.**
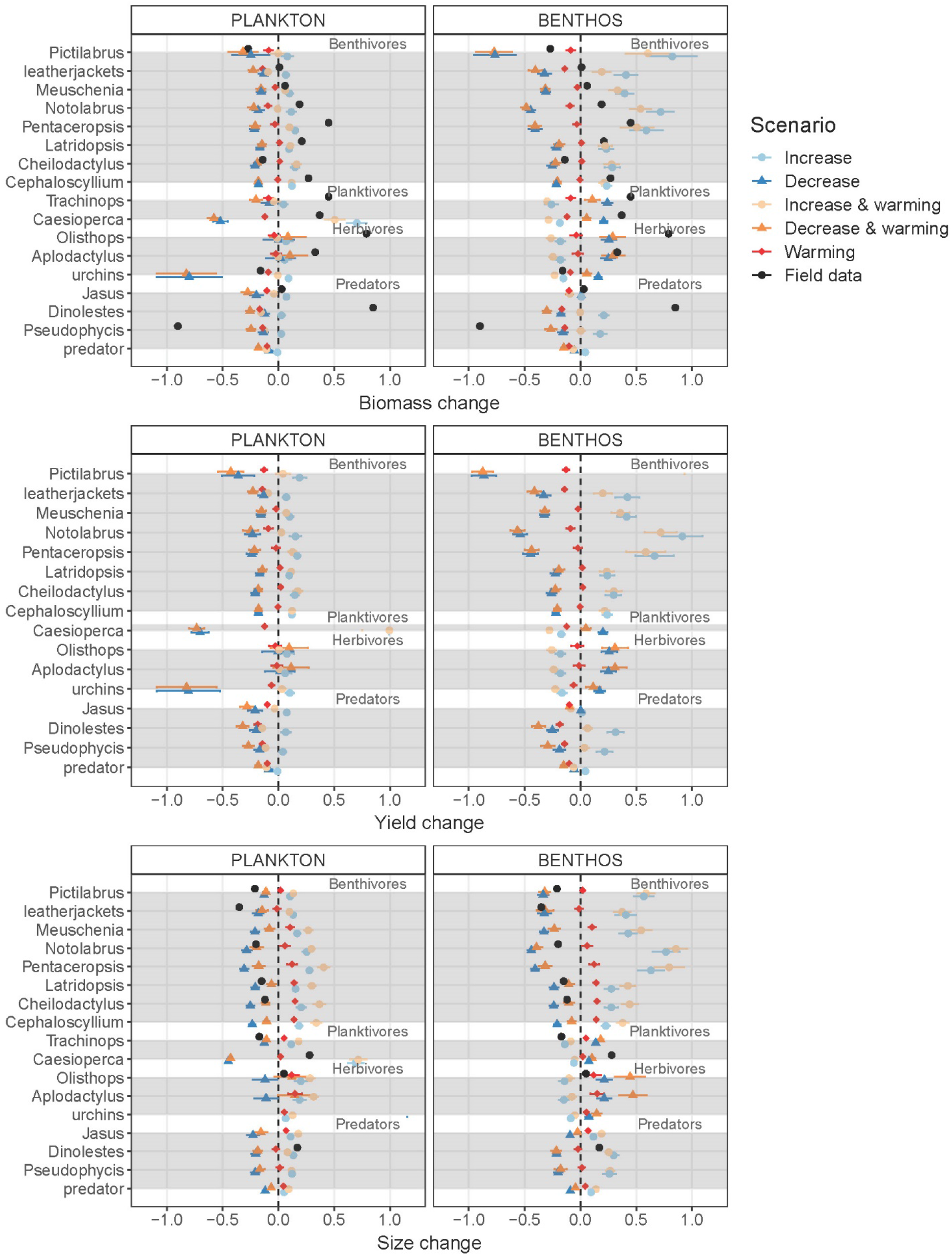
Changes in biomasses, yields and mean body sizes (individuals above 2cm length) in model groups across the alternative model scenarios with increasing or decreasing resource (plankton or benthos) abundance. Variation across scenarios in all 29 parameter combinations is shown with horizontal bars. Empirically observed changes are shown with black dots. Yields of *Pictilabrus* in scenarios with increased benthos abundance where ca 1.3 (with warming) and 1.8 (no warming) times higher and are not shown here due to scale.

In general, warming resulted in predicted changes in species’ biomass or yields that were smaller than those from changes in resource abundance (Fig. 3). Responses to temperature either decreased or did not affect biomasses much, and the biomass response varied across species and trophic groups, despite assuming identical temperature sensitivity parameters for all of them (i.e., increased search, maximum intake and metabolism rates, higher background and senescence mortality at higher temperatures). This indicates that emergent biomasses and yields were affected by species interactions, which modified warming driven physiological and mortality impacts. Changes in plankton resource abundance had similar qualitative effects on biomasses and yields of most species, where biomasses of most species increased by 5-10% when plankton *κ* increased by 30% and decreased by 10-20% when *κ* decreased by 30% (with high variation in some species, light and dark blue symbols in Fig. 3).

To assess model predictions from alternative scenarios against field observations, we looked at mean biomass changes of the model species between 1990s (used for model calibrations) and those from 2015-2020 from field survey data. Over this period, the observed biomass of most species either increased or remained similar. Qualitatively similar responses across trophic groups were more consistent with changes in plankton than benthic resource alone, but the magnitude was more consistent with changes in the benthos resource (but see Discussion for the limitation of such comparison).

In contrast, changes in benthic resource abundance had opposing effects —biomass and yield of all benthivores changed in the same direction as benthos abundance, whereas the opposite was true for planktivores and herbivores. For predatory species, higher abundance of benthos increased biomass and yields of *Dinolestes lewini* and *Pseudophycis palmata* by ca 20%, but had no effects on lobsters or the general predator. These contrasting responses across trophic groups were confirmed by the mixed effect ANOVAs (Fig. 4, Table S8), where species within a trophic group were treated as random effects and separate ANOVAs were performed for biomass, yield and size responses to plankton or benthos changes interacting with temperature change. The strongest response to changes in plankton was, as expected, seen for planktivore biomass and yields, whereas benthivores showed the strongest response to changes in benthic resource. An overall negative response of temperature was observed in biomass and yields of all four trophic groups, but the impact was strongest (and highly significant) in predators (red symbols in Fig. 4).

**Fig. 4.**
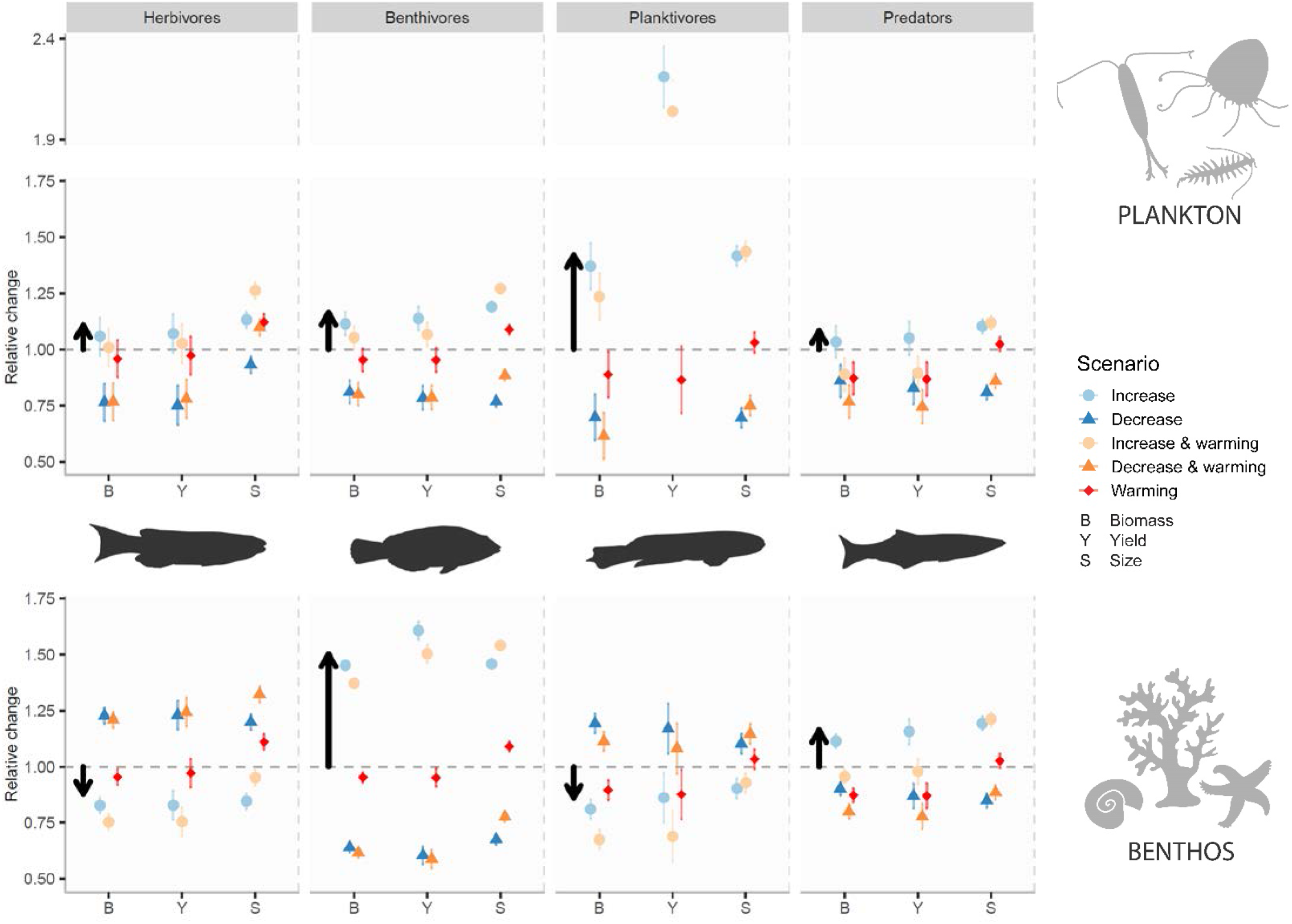
Mixed-effect ANOVA for predicted effects on biomass, mean size (for individuals larger than 2cm) and yield of four coastal reef fish trophic groups to changes in pelagic and benthic resource abundance and interaction with temperature, under low fishing scenarios. Error bars show 95% confidence intervals from ANOVA analyses. Black arrows emphasise the predicted effects of increased plankton or benthos abundance on biomass and mean size responses across trophic groups (same as light blue circles), and highlight their contrasting responses to changes in plankton versus benthos. Planktivore yield decreases due in reduced plankton scenarios was very strong (ca 0.25, below Y-axis minimum limit shown) and is not shown for clarity. For each parameter combination, all responses are scaled to values in baseline simulations (no resource change, no warming). Note, for planktivores biomass and size change reflects effects in two species, whereas yield change only shows responses in *Caesioperca* (as *Trachinops* is not fished).

When physiological responses to temperature were applied together with changes in resource levels, temperature usually had a slightly negative impact on species’ biomass, especially for scenarios where resource levels increased (light orange symbols in Figs 3, 4). This impact varied quantitatively across species, and was consistent with their responses to temperature impacts alone; i.e. species which had strongest biomass changes to temperature impacts alone (e.g. *D. lewini, P. palmata*) were also more negatively affected by temperature when this impact was combined with the resource change. At a trophic group level, any biomass and yield increase in predators due to higher benthos abundance were completely negated when increasing benthos levels were combined with higher temperatures (bottom right, Fig. 4).

### Higher temperatures increased mean body sizes of most species, but resource change had larger quantitative effect

As with biomass, the mean intra-specific body size change was considerably more influenced by resource changes than warming, and the relative magnitude of responses across species (10-30%) was equivalent to changes in biomass (Fig. 3). The largest changes in mean size (20-60% increase) were seen for benthivorous species under scenarios with increased benthos abundance. Unlike for biomass and yields, however, increases in temperature had a positive impact on mean size of most species, regardless of whether temperature increased independently of, or in combination with, the resource change. The increase in mean body size was strongest in herbivores and benthivores (ca 10%, red and orange symbols in Fig. 4). Strong positive effects of warming on mean body size were likely caused by higher food intake rates at higher temperatures, allowing for faster growth in early life. Across trophic groups plankton and benthos changes had similar qualitative effects on species mean sizes, as it had for biomass and yields. Namely, when plankton abundance increased, species’ mean sizes also increased (or remained similar). In contrast, when benthos abundance increased, only benthivorous and predatory species became larger, whereas species in planktivore and herbivore groups became slightly smaller (Figs 3, 4). Comparison of model predictions with observed temporal mean body size changes was possible for many of the benthivorous species using estimates from (Audzijonyte et al. 2020). Observed body size changes since the 1990s (black dots in Fig. 3C) were generally consistent in magnitude and direction with predictions from scenarios with 30% decreases in plankton or benthic resource abundance (dark orange and blue symbols in Fig. 3C).

## Discussion

### Changes in benthic resource abundance have major and contrasting impacts on different trophic groups of coastal fish

By introducing independent size-structured plankton and benthic production pathways and exploring how relative changes in their abundance affect coastal fish communities, our study showed that biomasses and mean sizes of different trophic groups can respond to benthic resource changes in qualitatively different ways. Increased abundance of benthic resources led to higher biomasses, yields and mean sizes of benthivores and predators, but the reverse was true for herbivores and planktivores. Such opposing responses emphasise the importance of species interactions and possible feedback loops (Audzijonyte et al. 2013), and suggest a potentially rapid reorganisation of fish communities and food webs, with yet unclear impacts on coastal fisheries. In contrast, changes in plankton abundance induced similar responses across all trophic groups, such that decreasing plankton abundance affected all species negatively. These findings are consistent with other regional and global ocean models, suggesting decreasing fish biomass in response to decreasing plankton primary productivity (Blanchard et al. 2012, Bryndum-Buchholz et al. 2019), and in line with the expectations of size-based multispecies models, where all species initially feed on the smallest sizes of plankton and are all impacted by changes in plankton.

Marine ecosystem climate change studies and models have been strongly focused on understanding changes in plankton primary production, which drives fish production in pelagic and shelf ecosystems and supports huge fisheries. Yet our study emphasises the critical importance of changes in the benthic production pathway for the shallow water reef ecosystems that are of extreme importance to coastal communities, not just for food production, but also tourism and recreation and numerous goods and services. While plankton productivity can be estimated, modelled and predicted over huge scales using remote sensing data, very little knowledge exists on how benthic energy pathways vary spatially, temporally or will respond to changing ocean climates. In addition to changing sea temperatures and nutrient profiles, coastal seas will also experience productivity changes resulting from climate driven precipitation changes on land, altered land-use and urbanisation. Moreover, benthic primary production can come from multiple sources, including large fleshy macroalgae (such as kelps), turfs with rapid turnover, and epiphytic algal production on all sunlit reef surfaces, each subjected to different ecological and environmental pressures. Predicting changes in these and understanding key drivers remain monumental scientific hurdles, emphasising the importance of considering alternative scenarios, as undertaken here.

The importance of integrating benthic organisms into community size spectra and ecosystem models is well recognised (Mehner et al. 2018) and supported by recent evidence suggesting that the size structure of coastal reef communities aligns better with theoretical expectations after integrating macroinvertebrates with fishes into one community size spectrum (rather than considering fishes in isolation, as traditionally done (Heather et al. 2021)). Multiple resource spectra have previously been included in marine ecosystem models, including by (Blanchard et al. 2010), who showed that coupled pelagic and benthic detrital resource pathways increases fish community stability. However, in that study, the benthic pathway was dependent on the pelagic production and not size structured. A benthic resource spectrum was also found essential for accurate representation of ontogenetic diet shifts in the Bering Sea multi-species size spectrum model (Reum et al. 2019b), but in that case the resource abundance was static (i.e. not dependent on the imposed mortality). End-to-end ecosystem models, such as Atlantis (Audzijonyte et al. 2019b) also introduce separate benthic resource groups (Bossier et al. 2018, Pethybridge et al. 2019), but they are modelled either as unstructured biomass pools or require detailed parameterisation to represent age-based growth dynamics. The modelling framework introduced in our study provides a straightforward and robust approach to integrate two or more independent and size structured energy pathways into modelling in a way that allows pooling benthic species into one general resource while preserving size structure and size-based feeding.

Preserving size structure of the benthic community is important because different sized benthic organisms can respond to climate change in opposing ways (Fraser et al. 2021). In our study, steepening resource size slopes and a resultant shift of resource abundance towards smaller organisms had an overall positive impact on fish biomass (Fig. S7). This is because most food limitation occurred at small fish body sizes, and large predator-prey mass ratios of benthivorous and planktivorous fishes (Coghlan et al. 2022) means that most fishes were feeding on smallest benthic size groups. An important future development for better representation of benthic resources in coastal marine ecosystems would be to link resource parameters (availability to predation, regeneration rate) to habitat complexity. For coral reef ecosystems, adding size-based predation refuges improved the fit of modelled fish size spectra to empirical observations (Rogers et al. 2014). For temperate rocky reefs, as studied here, habitat complexity is provided by large kelp stands in addition to rock crevices, and rapid loss of kelp forests is likely to reduce benthic productivity with major implications for temperate rocky reef communities and SE Tasmanian fisheries, which include benthivorous fishes (*Notolabrus*, banded morwong, trumpeter, leatherjackets).

### Physiological responses to temperature lead to lower biomass and larger mean sizes in individual species, but community level impacts are relatively small

Compared to changes in plankton or benthos abundance and size structure, the predicted physiological consequences of warming seas on biomass and mean size of fishes and large invertebrates were relatively small. The relative magnitude of these impacts varied across species and trophic groups, despite our model assuming that species’ shared the same temperature response parameters. Although we recognise that this assumption is unlikely, the results imply that in natural communities observed body size and biomass changes due to warming are strongly shaped by species interactions and therefore hard to predict *a priori*.In our study, physiological changes, both when analysed separately and in combination with resource change, had largest impacts on predator biomass and yields, and very little on herbivore and benthivore biomass (Fig. 4). For predators, such as *Dinolestes lewini* or commercially valuable and ecologically important rock lobsters, the negative physiological effect of increasing temperature negated any positive impacts on their biomass from increasing benthos abundance. This appears to be a food limitation response, in which these predators were unable to compensate increased metabolic needs with higher food intake. This finding is in line with other recent modelling studies (Lindmark et al. 2021) and empirical observations demonstrating that fish responses to warming are mediated by food availability.

A number of experimental and modelling studies have reported that warming waters will lead to smaller fish (Daufresne et al. 2009, Cheung et al. 2013, Pauly and Cheung 2018). At an intraspecific level this is believed to be driven by the thermal dependence of physiological processes (Angilletta Jr et al. 2004, Brown et al. 2004, Forster et al. 2011), although the exact mechanisms remain debated (Pauly and Cheung 2018, Audzijonyte et al. 2019a, Wootton et al. 2022). In contrast our study reports that physiological responses to warming generally increased, rather than decreased, mean species body sizes. This does not necessarily contradict the temperature-size rule or “shrinking” fish paradigm, because this paradigm at an intra-specific level refers to negative temperature impacts on maximum rather than mean ectotherm body sizes (Atkinson 1994). At the community level, the “shrinking fish” observation typically refers to redistributions from large bodied to small bodied species (Daufresne et al. 2009). Yet, impacts of warming on mean intra-specific body sizes are less clear. Since the majority of individuals in a population consist of juveniles, changes in mean and maximum body sizes may oppose each other, although testing this hypothesis requires further investigation. Our study shows that in a multi-species context, species interactions will modify both temperature and primary production impacts on fish biomass, yields and mean sizes, and is consistent with empirical analyses of temperature impacts on mean coastal fish body sizes (Audzijonyte et al. 2020), demonstrating varied mean body size responses across species.

Of course, conclusions from our study are highly dependent on our model assumptions and these assumptions are greatly simplified. There are at least three key assumptions that should be explored in further studies. The first relates to the magnitude and combination of resource and temperature responses. While we show that resource changes have greater impacts than temperature changes, clearly, this will depend on the relative magnitude of change assumed in the model. In our simulations, both productivity and temperature changes were deliberately high, yet within a realistic range (see Supplement). More specific scenarios were not possible due to high uncertainty in the expected global warming driven productivity and size structure changes, especially for the benthos (Barneche et al. 2021, Fraser et al. 2021). In reality, abundance and size structure of both plankton and benthic resources will change simultaneously, with different combination of temperature impacts. It was not feasible to address all possible resource and temperature change scenarios here, but to encourage further studies we provide an online ShinyR tool (https://fishsize.shinyapps.io/BenthicSizeSpectrum) to explore how combinations of resource abundance, size structure and temperature change will affect modelled species biomass, yields and mean body sizes.

Second, to keep the number of scenarios tractable, we assumed that all species and all vital rates (search rate, maximum food intake, metabolism, background and senescence mortality) scale with temperature in the same way. This is along the lines of metabolic theory of ecology (Brown et al. 2004), but in reality, temperature responses will differ across rates, species, ontogeny, populations and through time (due to acclimation and adaptation) (Barneche et al. 2021), (Pörtner 2002, Lindmark et al. 2018), (Donelson and Munday 2015, Denderen et al. 2020). Third, we assumed that temperature mostly impacts vital rates, but not life-history trade-offs such as relative energy allocation to growth and reproduction (Arendt 2011, Jorgensen et al. 2016, Ayllón et al. 2021). Some degree of growth-reproduction trade-off does occur in size-based models, because growth and therefore maturation (set to occur over some size range) are not set, but are emergent model properties (Andersen et al. 2007). Recent inter-generational experiments show that some populations may be capable of complete metabolic acclimation to higher temperatures, and that changes in growth and size may be largely driven by reproductive decisions (Wootton et al. 2022). Future studies should explore more direct impacts of temperature on these trade-offs by allowing decreasing maturation sizes, consistent with TSR observations and different reproductive allocation curves. All these modifications are easy to simulate in the current flexible *mizer* framework.

#### Future directions and limitations

Adequate prediction of coastal community responses to warming requires an understanding of interactive resource and physiological changes in multi-species contexts. Yet, marine ecosystem responses to climate change also involve other factors with potentially large impacts. The main one relates to the possible impacts of species redistributions (Pecl et al. 2014), where newly arriving species (e.g. sea urchins in kelp forest ecosystems) might cause major ecosystem shifts. Redistribution of herbivorous fish species into temperate reefs is already driving large changes in macroalgal abundance and community composition (Vergés et al. 2016). The second major one is human impacts, through either increased fishing or increased protection. An increasing number of studies shows that protecting large fishes, and especially predators, can help mitigate population and ecosystem impacts of warming (Bates et al. 2017, Wootton et al. 2021). Our study did not explore how different fishing scenarios might interact with global warming, but such analysis would be very important and relatively straightforward. More work should also be done to explore parameter uncertainty. While the current study presents one of the most exhaustive parameter uncertainty evaluations across a range of different traits (recruitment, food intake rates and species interactions), we didn’t explore the uncertainty in species specific physiological parameters, such as combination of maximum intake, search rate and metabolic constants, and how these might affect emergent species body sizes and abundance. Finally, the current framework also allows assessment of climate change impacts on broader community attributes and indicators used in ecosystem management and ecology. These include such characteristics as resilience and community connectedness, and ecological and fisheries indicators based on body size distributions in fish communities, overall community level size structure and maximum sustainable yield, or ecosystem level energy transfer efficiency. Our study provides a pathway that will hopefully encourage and facilitate further investigation.

## Acknowledgements

We thank Gretta Pecl and Jonathan Reum for suggestions for this study, Freddie Heather for help with the ShinyR tool, Amy Coghlan for model illustration (Fig. 2), Lara Beckmann for help with Figure 4, and many volunteers who helped with RLS surveys. This study was supported by the ARC Discovery grant DP170104240 and Pew Fellows Program in Marine Conservation. The ecological data used for this study are managed through, and were sourced from, Australia’s Integrated Marine Observing System (IMOS) – IMOS is enabled by the National Collaborative Research Infrastructure Strategy (NCRIS).

## Notes

### Competing Interest Statement

The authors have declared no competing interest.

