## Supplementary material for "Changes in benthic and pelagic production interact with warming to drive responses to climate change in a temperate coastal ecosystem"

#### Contents

1. Methods ... 3
1.1. Model assumptions and equations ... 3
1.2. Selection of species and functional groups ... 5
1.3. Model parameterisation ... 6
***Derivation of h*** ... 8
***Derivation of ks*** ... 10
1.4. Feeding interactions ... 11
***Interaction with resources*** ... 11
***Interaction among size structured groups*** ... 13
1.5. Calibrating reproduction parameters for realistic vulnerability to fishing ... 14
1.6. Model calibration procedure ... 15
***Initial parameter tuning*** ... 15
***Estimating parameter uncertainty using a rejection approach*** ... 18
1.7. Background resource parameters: plankton slope and abundance ... 20
Plankton slope ... 20
Plankton abundance ... 21
1.8. Background resource parameters: benthos slope and abundance ... 21
2. Results ... 23
2.1. Baseline biomass variation ... 23
2.2. Changes in the resource slopes and impacts on equilibrium biomasses ... 23
2.3. Mixed effect model selection and results ... 25
3. References ... 29

### 1. Methods

#### 1.1. Model assumptions and equations

Availability of food $E_{a,i}$ for each size group in a species *i* is dynamic and depends on the availability of all prey species *j* and background resources *R* and size preference function of the species *i*:

$$E_{a,i}(w_i) = \int \left( \sum_R \theta_{i,R} N_R(w_j) + \sum_j \theta_{i,j} N_j(w_j) \right) \phi_i\left(\frac{w_j}{w_i}\right) w_j dw_j \qquad \text{Eq1}$$

where $\theta_{i,j}$ is the availability of species *j* to species *i* (species interaction matrix, see below), $\theta_{i,R}$ is the availability of resource *R* for species *i* (where we use multiple resources, see below and Table 2), $w_j$ is the weight of prey, and $\phi_i\left(\frac{w_j}{w_i}\right)$ is the log-normal size selection function (Ursin 1967, Andersen 2019)

$$\phi_i\left(\frac{w_j}{w_i}\right) = \exp\left[\frac{-\left(ln\left(w_i/(w_j\beta_i)\right)\right)^2}{2\sigma_i^2}\right] \qquad \text{Eq2}$$

defined by the preferred predator-prey mass ratio of the species (predator) *i* ($\beta_i$) and the standard deviation of size selection function $\sigma_i$ (Table 2). The feeding kernel is truncated at a limit $w_i = w_j$, to ensure that consumers never feed on prey larger than themselves.

Energy flow in an individual follows general bioenergetic principles (see (Andersen 2019) ), where available food ($E_a$) is consumed based on the Holling type II feeding response (Eq 7, Tables S1, S2), assimilated based on assimilation efficiency $\alpha$ (Eq 9), used to cover maintenance costs that scale with body size (Eq 9) and is then allocated between growth and reproduction depending on the maturity status (Eq 10, 11). Proportion of energy allocated to reproduction assumes an asymmetric logistic function, with the midpoint determined by the maturation size ($w_m$), steepness parameter ($u$) and the rate at which allocation approaches 1 determined by the scaling exponent of the maximum intake rate (*n*) (Eq 10) (Andersen 2019).

**Table S 1.** Model equations.

| Description | Expression | # |
|---|---|---|
| Encountered food | $E_{e,i}(w) = \gamma_i w^q E_{a,i}(w)$ | Eq3 |
| Maximum consumption rate | $h_i w^n$ | Eq4 |
| Feeding level (Holling type II) | $f_i(w) = \dfrac{E_{e,i}(w)}{E_{e,i}(w) + h_i w^n}$ | Eq5 |
| Food consumption rate | $f_i(w) h_i w^n$ | Eq6 |
| Net energy | $E_{r,i}(w) = \max\left(0, \alpha f_i(w) h_i w^n - k_{s,i} w^p\right)$ | Eq7 |
| Proportion of energy invested into reproduction | $\psi_i(w) = \left[1 + \left(\dfrac{w}{w_{m,i}}\right)^{-u}\right]^{-1} \left(\dfrac{w}{w_{\infty,i}}\right)^{1-n}$ | Eq8 |
| Somatic growth | $g_i(w) = \alpha_g E_{r,i}(w)\left(1 - \psi_i(w)\right)$ | Eq9 |

| | | |
|---|---|---|
| Total egg production | $$R_{p,i} = \frac{\varepsilon}{2w_R}\int N_i(w)E_{r,i}(w)\psi_i(w)\,dw$$ | Eq10 |
| Recruitment | $$R_i = R_{max.i}\frac{R_{p,i}}{R_{p,i}+R_{max,i}}$$ | Eq11 |
| Background mortality | $$\mu_{b,i} = \mu_0 w_\infty^{-0.25}$$ | Eq12 |
| Starvation mortality | $$\mu_{s,i}(w) = \max\left(0,\frac{k_{s,i}w^p - \alpha f_i(w)h_i w^n}{\xi w}\right)$$ | Eq13 |
| Predation mortality | $$\mu_{p,j}(w_j) = \sum_i \int \phi_i\left(\frac{w_j}{w}\right)(1-f_i(w))\gamma_i w^q \theta_{ij} N_i(w)dw$$ | Eq14 |
| Senescence mortality | $$\mu_{sc,i}(w_i) = \vartheta\, 10^{\rho(ln(w)-\ln(x_s w_\infty))}$$ | Eq15 |
| Fishing mortality | $$\mu_{f,i}(w_i) = \begin{cases} 0, & w < w_{m,i} \\ F_i, & otherwise \end{cases}$$ | Eq16 |

**Table S 2.** General model parameters.

Species specific parameters are indicated as SS and shown in Tables S4 and S5. Parameters marked with * were explored in uncertainty analyses.

| Symbol | Description | Units | Value | Source |
|---|---|---|---|---|
| $w_R$ | Size at recruitment (smallest size) | g | SS, Eq 17 | after (Barneche et al. 2018) |
| $w_\infty$ | Maximum body size | g | SS | This study |
| $w_m$ | Body size at maturation | g | SS | This study |
| | ***Feeding and growth*** | | | |
| *n* | Exponent of max. consumption | - | 2/3 | Hartvig 2011 |
| *q* | Exponent of search volume | - | 0.8 | Hartvig 2011 |
| *p* | Exponent of standard metabolism | - | 0.7 | Hartvig 2011 |
| *h* | Constant of max. consumption | g yr$^{-1}$ g$^{-n}$ | SS, Eq 18 | This study |
| $\gamma$ | Constant of search volume | m$^2$ yr$^{-1}$ g$^{-q}$ | SS* | This study |
| $k_s$ | Constant of standard metabolism | g yr$^{-1}$ g$^{-p}$ | SS, Eq 19 | This study |
| $\alpha$ | Assimilation efficiency | - | 0.6 or 0.2 | (Hartvig et al. 2011) |
| $\alpha_g$ | Growth efficiency | - | 0.6 | This study |
| $\beta$ | Preferred predator-prey mass ratio | - | SS | This study |
| $\sigma$ | Width of the predation kernel | - | SS | This study |
| | ***Reproduction*** | | | |
| $\varepsilon$ | Reproductive efficiency | - | SS, tuned | |
| $R_{max}$ | Maximum recruitment | ind m$^{-2}$ | SS, tuned* | |
| *u* | Maturation transition width | - | 5 | |
| | ***Mortality*** | | | |
| $\mu_0$ | Background mortality constant | yr$^{-1}$ | 0.3 | Modified after Hartvig 2011 |
| $\xi$ | Starvation mortality constant[1] | yr$^{-1}$ | 0.1 | Hartvig 2011 |
| $\vartheta$ | Senescence mortality constant[2] | yr$^{-1}$ | 0.1 | Modified after (Law et al. 2009)) |
| $x_s$ | Proportion of $w_\infty$ at which sencescece mortality is equal to $\vartheta$ | - | 0.95 | As above |
| $\rho$ | Senescence mortality exponent | - | 3 | (Law et al. 2009) |

| | | | | |
|---|---|---|---|---|
| $S$ | Size based fisheries selectivity, here assumed to be knife edge at $w_{m,i}$ | - | | |
| $F_i$ | Instantaneous fisheries mortality of fully selected individuals | $yr^{-1}$ | SS | This study |
| | **Background resource spectra** | | | |
| $\lambda_P$ | Exponent of pelagic resource spectrum | - | Table 2 | This study |
| $\lambda_B$ | Exponent of benthic resource spectrum | - | Table 2 | This study |
| $\lambda_A$ | Exponent of the macroalgal spectrum | | 1.6 | This study |
| $\kappa_P$ | Pelagic resource spectrum constant | $g\ m^{-2}$ | Table 2 | This study |
| $\kappa_B$ | Pre-factor for benthic resource spectrum | $g\ m^{-2}$ | Table 2 | This study |
| $\kappa_A$ | Pre-factor for macroalgal resource spectrum | $g\ m^{-2}$ | 16 | This study |
| $r_{0P}$ | Relagic resource regeneration rate | $g\ m^{-2}$ | 1 | This study |
| $r_{0B}$ | Benthic resource regeneration rate | $g\ m^{-2}$ | 1 | This study |
| $r_{0A}$ | Macroalgal resource regeneration rate | $g\ m^{-2}$ | 2 | This study |
| $[w_{min}, w_{max}]_P$ | Size range of the pelagic resource spectrum | g | $1^{-10}$, 1 | This study |
| $[w_{min}, w_{max}]_B$ | Size range of the benthic resource spectrum | g | $1^{-3}$, 5 | This study |
| $[w_{min}, w_{max}]_A$ | Size range of the algal resource spectrum | g | $1^{-3}$, 50 | This study |

[1]- not included in standard MSSS (or in 'mizer'), but available through *mizerStarvation* extension and used in this study

[2]- senescence mortality not applied to species with the maximum body size < 400g

#### 1.2. Selection of species and functional groups

A successful calibration and performance of a multi-species models requires that groups included in the model can be differentiated using their modelled traits, such as maximum size, maturation size, life-history characteristics or diets. Further, increasing the number of species leads to increase in model complexity and parameter numbers. We therefore aimed to find a balance between model simplicity and realistic representation of the system relevant to the question (benthic and pelagic food web dynamics).

To select species to be included in the model we used underwater visual survey data from Australian Temperate Reef Collaboration (ATRC) monitoring program (Edgar and Barrett 2012). The model was calibrated using survey data the 1990s from the coasts of Tasmania, excluding the biogeographically distinct northern Tasmanian coast. This data set included 1965 underwater method 1 surveys ($50m^2$) (for further details on survey methodology see (Edgar et al. 2016)). We selected all species that occurred in more than 2.5% of surveys and had the average biomass per survey of at least $25g/50m^2$, and removed 7 rare species that had low abundance and do not play a critical role in the ecosystem (e.g. are not predators) (*Girella zebra, Cheilodactylus nigripes, Diodon nicthemerus, Caesioperca lepidoptera, Pseudolabrus mortonii, Urolophus cruciatus, Neoodax balteatus*) and pooled species with similar life-history and diets. Specifically, the two common *Notolabrus* species have similar life-history characteristics and diet and have been pooled into one group. Further, *M. australis* and *A. vittiger* both grow to about 320mm, have similar growth trajectories (they reach about 15-18cm by the age of 2, and 30 cm by the age of 5-6 and older ages are not seen), and feed on invertebrates, such as molluscs, echinoids and poriferans (Barrett 1995). They are pooled into a leatherjacket category. This procedure gave 14 fish species/groups. In addition we added two invertebrate groups (lobsters and urchins) in the size structured group list (see main manuscript).

**Table S 3.** Final list of model species and groups, with the common names, scientific names and model abbreviation names.

$L_\infty$ are the maximum lengths, $w_\infty$ and $w_m$ are maximum and maturation body size (in grams), biomass (g/m$^2$) is equilibrium biomass from surveys from 1992 to 2000, used to parameterise $R_{max}$ values.

| Common name | Scientific name | Model name | $L_\infty$ | $w_\infty$ | $w_m$ | Biomass (g/m$^2$) |
|---|---|---|---|---|---|---|
| Purple wrasse and bluethroat wrasse | *Notolabrus tetricus & N. fucicola* | Notolabrus | 40-50 | 1600 | 250 | 2.040 |
| Bastard Trumpeter | *Latridopsis forsteri* | L_forsteri | 62 | 5193 | 792 | 0.653 |
| Hulafish | *Trachinops caudimaculatus* | T_caudimaculatus | 15 | 39 | 7 | 0.580 |
| Barber Perch | *Caesioperca rasor* | C_rasor | 25 | 623 | 161 | 0.749 |
| Long-fin Pike | *Dinolestes lewini* | D_lewini | 50 | 1032 | 250 | 0.327 |
| Leatherjackets | *Acanthaluteres vittiger & Meuschenia australis* | leatherjackets | 40 | 2974 | 650 | 0.776 |
| Banded Morwong | *Cheilodactylus spectabilis* | C_spectabilis | 87.5 | 13568 | 3500 | 1.008 |
| Draughtboard Shark | *Cephaloscyllium laticeps* | C_laticeps | 150 | 16012 | 2100 | 0.187 |
| Senator Wrasse | *Pictilabrus laticlavius* | P_laticlavius | 30 | 389 | 51 | 0.404 |
| Long-snouted Boarfish | *Pentaceropsis recurvirostris* | Boarfish | 50 | 2192 | 275 | 0.058 |
| Six-spine Leatherjacket | *Meuschenia freycineti* | M_freycineti | 55 | 4300 | 1012 | 0.126 |
| Red Cod | *Pseudophycis palmata (former P. bachus)* | P_bachus | 50 | 1512 | 325 | 0.080 |
| Herring Cale | *Olisthops cyanomelas* | O_cyanomelas | 63 | 3364 | 700 | 0.099 |
| Marblefish | *Aplodactylus arctidens* | A_arctidens | 75 | 4400 | 803 | 0.220 |
| Predators | | predator | | 5000 | 800 | 1* |
| Urchins | *Heliocidaris, Centrostephanus & Goniocidaris* | urchins | | 350 | 50 | 2.642 |
| Rock lobsters | *Jasus edwarsii* | lobsters | | 3000 | 500 | 1.380 |

* - the biomass of predators is not taken from surveys, but assumed here to represent potential predation from all large predators (birds, marine mammals, mobile large sharks and pelagic predators)

##### 1.3. Model parameterisation

General model parameters, such as scaling exponents and mortality parameters were taken from earlier size spectrum model applications (Table 2 and references therein). We assumed standard food assimilation efficiency (α) for most model groups, but lower values (0.2) for the three herbivore species. Growth conversion efficiency or cost of growth is not typically included in MSS models (more specifically, it is accounted for by the general food assimilation efficiency α), but is introduced here (Eq11) to account for the cost of growth in line with dynamic energy budget models (Kooijman and Kooijman 2010) or optimal allocation models (Jørgensen and Fiksen 2010) (Audzijonyte and Richards 2018). The cost of growth ($\alpha_g$) is assumed to be at 0.6 as an average of both reserve, gonad and structure growth used in models above. Following (Hartvig et al. 2011) background mortality is scaled with species maximum body size, leading to species-specific but size independent (within species) background mortality rates (Table 3). However, we used a twice lower mortality constant (0.3 instead of 0.6) (Eq12) to reduce constant non dynamic background mortality and give more weight to dynamic mortality terms that emerge from species and resource interactions (predation, starvation).

Maximum species body sizes (in length) were extracted from the maximum observed body sizes in the surveys and then modified according to the expert advice (Rick Stuart-Smith, Neville Barrett and Graham Edgar). Maturation sizes were not available for all species, and in order to follow general life-history invariants, maturation length was assumed to be at around 0.5-0.6 of maximum body length (Charnov et al. 2013); adjusted based on the expert opinion. The maturation and maximum length were then converted to weights based on weight-length conversion parameters available for the Reel Life Survey species (Stuart-Smith et al. 2013). We used these general relationships to represent general coastal species of different body sizes and trophic roles, rather than specifically parameterise them to the Tasmanian coastal system.

Minimum size was calculated according to (Barneche et al. 2018) who calculated egg sizes across a range of female sizes. The smallest size class in the model (recruits) was estimated a 5x egg size to reduce the number of smallest size groups for computational efficiency and because dynamics in smallest size groups is anyway not reflected accurately in a spatially unstructured and deterministic model.

$$w_{R,i} = \left(0.0015 w_{\infty,i}^{0.14}\right) * 5 \qquad \text{Eq17}$$

The original (Barneche et al. 2018) relationship was applied to both intra and inter-specific relationships and used spawning female size rather than the maximum species body size. However, given that the maturation in this study is set as a proportion of maximum body size, the spawner size is correlated to the maximum body size (largest species will on average have larger spawners). The assumed minimum size is only an approximation, but unlikely to have large effects on egg production or early mortality. This is because the total egg production is determined by egg size and reproduction efficiency (Eq 10) and reproduction efficiency is calibrated for different species to give reasonable sensitivity to fishing (see below); predation mortality meanwhile is similar across all smallest size groups.

Parameter values for the mass-specific search volume (γ), maximum intake rate (*h*) and metabolic constant (ks) were species specific. In earlier applications of MSS and by default in *mizer* species specific *h* values (g/g$^n$/year) are estimated from Von Bertalanffy growth parameters, feeding level and assimilation rates (Blanchard et al. 2004). However, for most species used in this study estimates of Von Bertalanffy growth rate coefficient were highly uncertain or not available. To get species-specific estimates of *h* we therefore derived an approximation of *h* from the maximum body size $(w_\infty, g)$, using daily size-specific intake rates estimates for a range of Australian coastal fish species (Soler et al. 2018) (see below for details on the "Derivation of h"):

$$h_i = 50\left(\frac{w_{\infty,i}}{1000}\right)^{0.15} \qquad \text{Eq18}$$

For herbivore species *h* values estimated from the equation above were increased 3 times to account for three times lower assimilation efficiency (α) (Table 2). The positive relationship between *h* and $w_\infty$, assumed above, means that we had to reconsider default MSSS assumption about standard maintenance cost *ks* (metabolic rate). By default, when n = p, in *mizer ks* = 0.12*h*, assuming that intake equals metabolism at the feeding level of 0.2 (critical feeding level) and that assimilation efficiency is 0.6 (0.6x0.2 = 0.12 gives the proportion of maximum intake required to cover the maintenance cost). However, since *h* scales positively with $w_\infty$ (Eq18), this would also mean positive relationship between *ks* and $w_\infty$. The observed "cost of life" or mass-specific metabolic rates at same body sizes are often higher is species that grow to smaller maximum body sizes (Kooijman and Kooijman 2010) (see below). We therefore developed another approximation of species-specific standard metabolic rate $(ks_i)$to maximum body size using DEB database values of the maintenance costs (see below for details on the "Derivation of *ks*"):

$$ks_i = 20\, w_{\infty,i}^{\ -0.25} \qquad \text{Eq 19}$$

**Table S 4.** Species specific parameters (see Table 2 for details).

Feeding parameters β and σ define the predation kernel, aP, aB and aA are vectors defining proportion of the pelagic, benthic and macroalgal resource spectra available for a given species. Parameters marked with * are defined using general equations (see above), parameters marked in italics are tuned to ensure emergent dynamics satisfy model criteria, and parameters marked in bold are further explored using uncertainty analysis.

| ***Species*** | ***Food selection*** | | | | | | ***Reproduction*** | | ***Intake and metabolism*** | | | ***Mortality*** |
|---|---|---|---|---|---|---|---|---|---|---|---|---|
| | β | σ | aP | aB | aA | **Trophic group** | $R_{max}$ | $\varepsilon$ | $\gamma$ | $k_s$* | $h$* | $\mu_0$* |
| Notolabrus | 500 | 1.3 | 0.2 | 0.7 | 0 | benthivore | **0.44324** | *0.3638* | **5.82** | 3.16 | 53.65 | 0.047 |
| L_forsteri | 2000 | 1.5 | 0.2 | 0.7 | 0 | benthivore | **0.00256** | *0.067* | **7.7** | 2.36 | 64.02 | 0.035 |
| T_caudimaculatus | 500 | 1.5 | 0.8 | 0 | 0 | planktivore | **0.88** | *0.4179* | **2.89** | 4.5 | 30.73 | 0.12 |
| C_rasor | 1000 | 1.5 | 0.8 | 0 | 0 | planktivore | **0.1755** | *0.1787* | **3.95** | 4 | 46.57 | 0.06 |
| D_lewini | 100 | 1.3 | 0.15 | 0.25 | 0 | predator | **0.09817** | *0.2034* | **20.82** | 3.53 | 50.24 | 0.053 |
| leatherjack | 500 | 1.3 | 0.2 | 0.7 | 0.02 | benthivore | **0.01641** | *0.1751* | **7.03** | 2.71 | 58.88 | 0.041 |
| C_spectabilis | 3000 | 1.5 | 0.2 | 0.7 | 0 | benthivore | **0.00127** | *0.0026* | **6.01** | 1.85 | 73.93 | 0.028 |
| C_laticeps | 3000 | 1.5 | 0.2 | 0.7 | 0 | benthivore | **0.00014** | *0.0525* | **6.65** | 1.78 | 75.79 | 0.027 |
| P_laticlavius | 250 | 1.5 | 0.2 | 0.7 | 0 | benthivore | **0.36468** | *0.2801* | **3.27** | 4.5 | 43.4 | 0.068 |
| Boarfish | 1000 | 1.3 | 0.2 | 0.7 | 0 | benthivore | **0.00593** | *0.1294* | **6.1** | 2.92 | 56.25 | 0.044 |
| M_freycineti | 1000 | 1.5 | 0.2 | 0.7 | 0.02 | benthivore | **0.00134** | *0.1551* | **5.8** | 2.47 | 62.23 | 0.037 |
| P_bachus | 100 | 1.3 | 0.15 | 0.25 | 0 | predator | **0.00849** | *0.1138* | **22.04** | 3.21 | 53.2 | 0.048 |
| O_cyanomelas | 500 | 3 | 0.2 | 0 | 0.7 | herbivore | **0.00658** | *0.1566* | **7.9** | 2.63 | 119.96 | 0.039 |
| A_arctidens | 500 | 3 | 0.2 | 0 | 0.7 | herbivore | **0.00703** | *0.1914* | **8.22** | 2.46 | 124.89 | 0.037 |
| predator | 100 | 1.3 | 0.15 | 0.25 | 0 | predator | **0.00319** | *0.0083* | **30.29** | 2.38 | 63.65 | 0.036 |
| urchins | 50 | 1.5 | 0.2 | 0 | 0.7 | herbivore | **0.74452** | *0.1155* | **11.35** | 4.62 | 85.43 | 0.069 |
| lobsters | 100 | 1.3 | 0.15 | 0.25 | 0 | predator | **0.0162** | *0.2097* | **26.87** | 2.7 | 58.96 | 0.041 |

#### *Derivation of h*

The goal of this approximation is to suggest a general relationship between $h_i$ and $w_\infty$ for species for which Von Bertalanffy growth parameters are not available and standard *mizer* approach cannot be used. For this purpose we estimated maximum intake rate constant for each model species using individual level species and weight specific daily intake rate values ($\text{DailyIntake}_{W,i}$) from (Soler et al. 2018). These intake values were estimated based on the empirically derived equation in (Palomares and Pauly 1989) and a statistical method to estimate diet composition (Soler et al. 2016). These daily intake rates range from 5-8% of body weight in small individuals to 0.5-1% in large individuals and are consistent with those used in experimental conditions to achieve realistic growth rates (Kjesbu et al. 1996, Skjæraasen et al. 2009); for herbivore species the daily intake values are several times larger. These daily intakes reflect the realised intake in wild conditions and not the maximum feeding rates. To translate the realised daily intake values to maximum daily intake rates we have to assume a certain feeding level.

First, we assume the default *mizer* feeding level (*f*) of 0.6 across all sizes, to get the maximum daily intake across sizes and species, where $\text{MaxDailyIntake}_{W,i} = \text{DailyIntake}_{W,i}/f$.

To get from individual level $MaxDailyIntake_{W,i}$ values to mass specific maximum intake rate and body size scaling exponent, we can apply a log-log linear regression of maximum daily intake (in grams) against body weight (in grams) for each model species *i* (see R code for further details).

log(MaxDailyIntake) = $a$ + $b$*log($W_{inf}$) Eq 20

The estimated maximum intake rate coefficient (which in the Eq20 corresponds to $e^a$) ranged around 0.10-0.11 g/$g^n$/day for non-herbivore species, while the body size scaling exponent (*n*, which in the equation above corresponds to *b*) was ~0.8. However, given that larger individuals typically have higher reserve densities than small individuals, feeding level under normal conditions could be expected to increase with body size. We therefore assumed that *f* increases smoothly with body as such as

$f(w) = f_{w\_min} + (f_{w\_max} - f_{w\_min})*e^{-z(Winf-w)}$, Eq 21

where $f_{w\_min}$ and $f_{w\_max}$ are feeding level values at the smallest and largest body size ($W_{inf}$), and z is the exponent defining how rapidly feeding level changes (here assumed to be 0.15 to give a smooth transition). We assumed that $f_{w\_min}$ = 0.5 and $f_{w\_max}$ = 0.9 values, derived new individual level maximum intake values from the daily intake values given in Soler et al. (2016), and again fitted a linear regression described above to estimated species-specific maximum intake versus body size coefficients, but this time using weight specific feeding levels, rather than a fixed feeding level of 0.6. These estimates gave an average maximum intake coefficient of 0.23 g/$g^n$ /day and the average body size scaling exponent *n* around 0.7. The exponent value was close to the n = 2/3 used in this model or DEB theory (Kooijman and Kooijman 2010), but the coefficient of 0.23 g/$g^n$/day was on the upper range of the values estimated in DEB database (0.06-0.19 g cm-2 day-1, DEB online database, Kooijman and Lika, 2014) or those used in the North Sea multispecies model (0.04-0.17 g/$g^n$/day, sizespectrum.org). Results were similar if $f_{w\_min}$ = 0.5 and $f_{w\_max}$ = 0.8 or if $f_{w\_min}$ = 0.6 and $f_{w\_max}$ = 0.9, but smaller differences between $f_{w\_min}$ and $f_{w\_max}$ gave higher body size scaling exponents and lower mass-specific constant.

The individual species maximum intake coefficient values estimated using the $f_{w\_min}$ = 0.5 and $f_{w\_max}$ = 0.9 assumption were then used to derive a general across species relationship between maximum intake coefficient and $w_\infty$ to be used as a general guidance for setting $h_i$ values in *mizer*. This was done by fitting another linear regression, this time between species specific maximum intake (now denoted as $h_i$ ) and $w_\infty$

$\log(h_i) = a + b*\log\left(\frac{w_{\infty,i}}{1000}\right)$, Eq 22

The estimated scaling exponent *b* was ~0.15 and exp(*a*) ~ 0.2. The value of 0.2 (estimated using maximum daily intake of g/g/day) was then converted to the annual maximum intake (g/$g^n$ /year) used in *mizer* (0.2*365 = 73), yielding a relationship of

$h_i = 73\left(\frac{w_{\infty,i}}{1000}\right)^{0.15}$ Eq 23

Applying this generic cross-species equation gives the $h_i$ values for the model species in the range of 70-90g/$g^n$/year. This is higher than defaults assumed in standard *mizer* trait model *h* = 40 g/$g^n$/year (Hartvig et al. 2011), North Sea multispecies model (18-61 g/$g^n$/year) (sizespectrum.org), DEB estimates of 20-73 g/$g^n$/year (DEB database) or ca 40 g/$g^n$/year estimates in optimal allocation models (e.g. Audzijonyte & Richards, 2019). This high estimate is likely to be due to the large number of herbivorous or partly herbivorous species in the (Soler et al. 2016) dataset. To correct for the high intake rates in this data set we used a slightly lower value scaling coefficient

$$h_i = 50\left(\frac{w_{\infty,i}}{1000}\right)^{0.15} \qquad \text{Eq 24}$$

This approximation is necessarily only a very crude one, but aims to account for the fact that, across datasets, larger bodied species often have higher mass-specific maximum intake rate values enabling early fast growth.

#### *Derivation of ks*

By default in *mizer*, when n = p, $k_s$= 0.12*h*. This is based on the assumption that critical feeding level is 0.2 and that assimilation efficiency is 0.6, which means that at 0.12*h* level intake just covers metabolism. If *h* ~ 40g/g/year (see above) this would give $k_s$ values of 4.8 g/g/year. Moreover, since *h* increases with maximum body size (Eq24), $k_s$ would also be larger in large-bodied fish. However, critical feeding level in small and large bodied species is unlikely to be similar, since "cost of life" is often higher is small bodied species. To attempt to estimate $k_s$ parameter that is independent of assumptions about *h* we extracted the DEB online database (https://www.bio.vu.nl/thb/deb/deblab/add_my_pet/ accessed on October 2018), where mass specific maintenance costs of structure are estimated for over 100 fish species of different body sizes. The average mass specific maintenance cost of structure at 20°C temperature in DEB is 20 J cm-3 day-1, but in slow growing vertebrates it can be as low as 10 J cm-3 day-1 (Kooijman 2000). The latter translates to 0.003 g/g/day or 1.1 g/g/year assuming 1g of structure mass equals 3000J and 1cm$^3$ of wet weight is 1g (van der Veer et al. 2009). Note, that this maintenance rate only applies to the cost of structure and, according to DEB assumptions, scales linearly with the structural mass (but not total mass). These values give an emergent total maintenance cost of an adult individual at 40-70% of its daily energy intake.

To derive a general relationship on how maintenance coefficients scales with maximum body size we fitted a linear regression to the DEB database columns p.M. (volume specific somatic maintenance rate) and zoom factor (maximum structural body length in cm or approximately $L_{max}$)

$\log(p.M_i) = a + b*\log(Zoom_i)$ Eq 25

The estimated intercept and slope of $p.M_i = 30.1 * L_{max,i}$^(-0.31) (Fig. S1). However, these estimates cannot be directly used in *mizer* due to different assumptions about scaling exponents and because DEB maintenance applies only to structural weight and not total body weight.

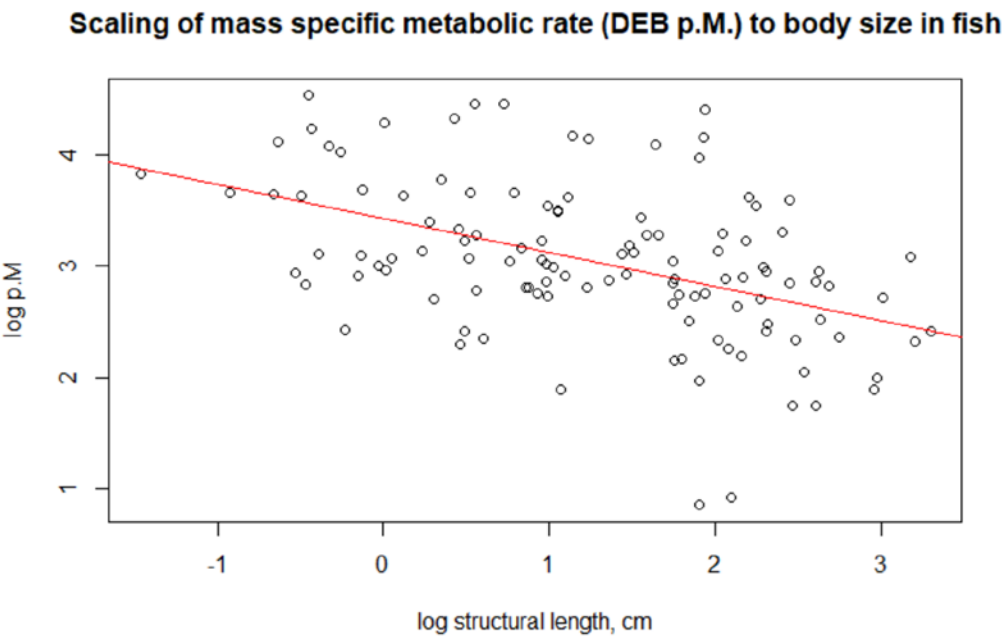

**Figure S 1.** Scaling of metabolism with body length in the Dynamic Energy Budget dataset.

Nevertheless, the data shows that metabolic rate coefficients ($k_s$) scales negatively with body size, and to follow the general fractal scaling rules of 0.25 or 0.75 we assume a negative scaling exponent of -0.25. The intercept was adjusted to yield $k_s$ values in the range of 3-5 g/g/year, used in the North Sea model (sizespectrum.org), giving the final general relationship

$$ks_i = 20\ w_{\infty,i}{}^{-0.25} \qquad \text{Eq 26}$$

For the smallest two species (urchins and hulafish) this equation gave too high $k_s$ values and they were reduced to 4.5 g/g$^p$/year (note the mizer units include -*p* because in mizer metabolism scales with body weight to the power of *p*, while in DEB scaling exponent of 1 to structural body mass is used).

#### 1.4. Feeding interactions

##### *Interaction with resources*

Availability of the three background spectra to each species are given in Table S5. All species have some access to the plankton spectrum, representing feeding in early life stages, but only planktivores have availability higher than 0.2. For predatory species access to the benthic spectrum is set at 25% to ensure ontogenetic diet switching from plankton, to benthic spectrum and then feeding on fish. Benthivorous species can access 70% of the benthic spectrum. The macroalgal spectrum is available only to herbivores, and a small proportion of it (2%) is available to leatherjacket species which are known to have algae in their diets (Soler et al. 2016). This formulation of the background spectrum availability, combined with different minimum and maximum sizes of the spectra, led to the emergent diet change from mostly pelagic food at larval stages, to benthos and, in species known to predate on other fish, to other fish (Fig. S2).

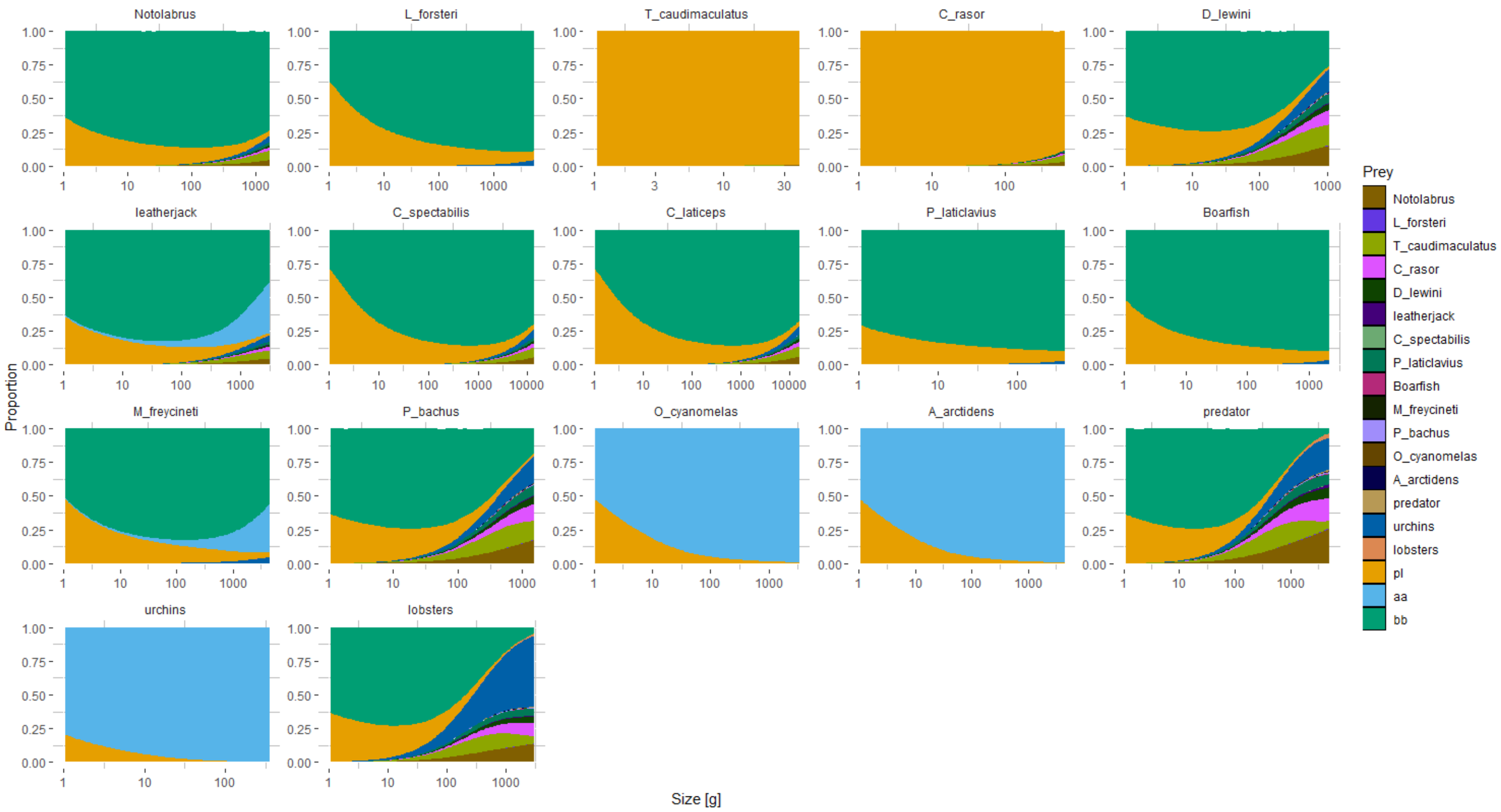

**Figure S 2.** Emergent species diets in baseline simulations.

For example, although all model groups have some access to the plankton spectrum, if availability of plankton spectrum is low, larger sizes switch to benthic spectrum (extending to larger sizes and having a shallower size spectrum slope), and then, depending on the species interaction parameters, to predation on fish. This formulation of model parameters produced emergent diets that were generally compatible with the available knowledge about species diets and were achieved with a small number of species-specific parameters. In Fig. S2 planktivorous species T_caudimaculatus and C_rasor mostly feed on plankton, although the larger C_rasor can consume small amount of small fish (larvae). Herbivorous species (O_cyanomelas, A_arctidens and urchins) transition from plankton to herbivory. Adult predators (D_lewini, P_bachus, predator, lobsters) mostly feed on other fish, but start by feeding on plankton and then on the benthic spectrum. The remaining benthivore species mostly remain feeding largely on the benthic resource, but can consume small amounts of small fish in their largest sizes. Up to now, such ontogenetic diet shifts have been difficult to replicate in physiologically structured multi-species models, without adding species details parameterised on empirical diet data (Bossier et al. 2018, Reum et al. 2019).

##### *Interaction among size structured groups*

Predatory interactions among size structured groups are represented by the interaction matrix, that can contain up to 17x17 = 289 parameters (Table S6). Here species interaction parameters are grouped into five categories:

*1)* 0 in cases where no interaction is allowed, i.e. herbivorous or strict benthivores fish don't feed on other fish,
2) 0.7 for all other cases where some predation is possible. Setting the value to 0.7 assumes that all prey biomass is never available to a predator at a given time (it still can be consumed over consecutive time steps).

The remaining three parameters were iteratively tuned to achieve species co-existence (Table S6) and later explored using uncertainty analyses (see below and Fig. S4). These parameters define

3) availability of small schooling pelagic fish to predators, assumed to be lower than for solitary fish due to predator avoidance schooling strategy. The tuned initial value was set at 0.4, alternative values in 29 parameter sets are shown in Fig. S4.
4) availability of urchins to fish, assumed to be lower due to morphological defences against predation, tuned to 0.15 (and Fig. S4 for the full range),
5) availability of urchins to lobsters, assumed to be higher than to fish, because lobsters can consume spiny urchins and urchins are an important part of lobster diet, tuned to 0.55 (and Fig. S4 for the full range, and Fig. S2 for the emergent lobster diet).

**Table S 5.** Availability of each species for predation by other species (interaction matrix).

Predators are in rows, prey are in columns. Values highlighted in bold were explored using the uncertainty evaluation (see Fig. S4 for full ranges).

| | Notolabrus | L_forsteri | T_caudimaculatus | C_rasor | D_lewini | leatherjack | C_spectabilis | C_laticeps | P_laticlavius | Boarfish | M_freycineti | P_bachus | O_cyanomelas | A_arctidens | predator | urchins | lobsters |
|---|---|---|---|---|---|---|---|---|---|---|---|---|---|---|---|---|---|
| Notolabrus | 0.7 | 0.7 | 0.4 | 0.4 | 0.7 | 0.7 | 0.7 | 0.7 | **0.4** | 0.7 | 0.7 | 0.7 | 0.7 | 0.7 | 0.7 | **0.15** | 0.7 |
| L_forsteri | 0 | 0 | 0 | 0 | 0 | 0 | 0 | 0 | **0** | 0 | 0 | 0 | 0 | 0 | 0 | **0.15** | 0.7 |
| T_caudimaculatus | 0.7 | 0.7 | 0.4 | 0.4 | 0.7 | 0.7 | 0.7 | 0.7 | **0.4** | 0.7 | 0.7 | 0.7 | 0.7 | 0.7 | 0.7 | 0 | 0 |
| C_rasor | 0.7 | 0.7 | 0.4 | 0.4 | 0.7 | 0.7 | 0.7 | 0.7 | **0.4** | 0.7 | 0.7 | 0.7 | 0.7 | 0.7 | 0.7 | 0 | 0 |
| D_lewini | 0.7 | 0.7 | 0.4 | 0.4 | 0.7 | 0.7 | 0.7 | 0.7 | **0.4** | 0.7 | 0.7 | 0.7 | 0.7 | 0.7 | 0.7 | **0.15** | 0.7 |
| leatherjack | 0.7 | 0.7 | 0.4 | 0.4 | 0.7 | 0.7 | 0.7 | 0.7 | **0.4** | 0.7 | 0.7 | 0.7 | 0.7 | 0.7 | 0.7 | **0.15** | 0.7 |
| C_spectabilis | 0.7 | 0.7 | 0.4 | 0.4 | 0.7 | 0.7 | 0.7 | 0.7 | **0.4** | 0.7 | 0.7 | 0.7 | 0.7 | 0.7 | 0.7 | **0.15** | 0.7 |
| C_laticeps | 0.7 | 0.7 | 0.4 | 0.4 | 0.7 | 0.7 | 0.7 | 0.7 | **0.4** | 0.7 | 0.7 | 0.7 | 0.7 | 0.7 | 0.7 | **0.15** | 0.7 |

| | | | | | | | | | | | | | | | | |
|---|---|---|---|---|---|---|---|---|---|---|---|---|---|---|---|---|---|
| P_laticlavius | 0 | 0 | 0 | 0 | 0 | 0 | 0 | 0 | **0** | 0 | 0 | 0 | 0 | 0 | 0 | **0.15** | 0.7 |
| Boarfish | 0 | 0 | 0 | 0 | 0 | 0 | 0 | 0 | **0** | 0 | 0 | 0 | 0 | 0 | 0 | **0.15** | 0.7 |
| M_freycineti | 0 | 0 | 0 | 0 | 0 | 0 | 0 | 0 | **0** | 0 | 0 | 0 | 0 | 0 | 0 | **0.15** | 0.7 |
| P_bachus | 0.7 | 0.7 | 0.4 | 0.4 | 0.7 | 0.7 | 0.7 | 0.7 | **0.4** | 0.7 | 0.7 | 0.7 | 0.7 | 0.7 | 0.7 | **0.15** | 0.7 |
| O_cyanomelas | 0 | 0 | 0 | 0 | 0 | 0 | 0 | 0 | **0** | 0 | 0 | 0 | 0 | 0 | 0 | 0 | 0 |
| A_arctidens | 0 | 0 | 0 | 0 | 0 | 0 | 0 | 0 | **0** | 0 | 0 | 0 | 0 | 0 | 0 | 0 | 0 |
| predator | 0.7 | 0.7 | 0.4 | 0.4 | 0.7 | 0.7 | 0.7 | 0.7 | **0.4** | 0.7 | 0.7 | 0.7 | 0.7 | 0.7 | 0.7 | **0.15** | 0.7 |
| urchins | 0 | 0 | 0 | 0 | 0 | 0 | 0 | 0 | **0** | 0 | 0 | 0 | 0 | 0 | 0 | 0 | 0 |
| lobsters | 0.7 | 0.7 | 0.4 | 0.4 | 0.7 | 0.7 | 0.7 | 0.7 | **0.4** | 0.7 | 0.7 | 0.7 | 0.7 | 0.7 | 0.7 | **0.55** | 0.7 |

#### 1.5. Calibrating reproduction parameters for realistic vulnerability to fishing

Assumptions about the stock-recruitment relationship in each species is likely to have large effects on the emergent species response to fishing. This response will be determined by species reproduction parameters, such as maturation size, reproduction allocation curve, as well as reproduction efficiency and maximum recruitment parameters ($\varepsilon$ and $R_{max}$) (see (Andersen 2019), chapter 4, (Jacobsen et al. 2017) for further details). Therefore, in order to achieve reasonable species dynamics in response to fishing it is important to ensure that linear and non-linear terms of recruitment dynamics (defined by $\varepsilon$ and $R_{max}$) are in a reasonable range.

Because both reproduction efficiency ($\varepsilon$) and maximum recruitment ($R_{max}$) combine multiple real life processes, they individually cannot be calibrated to observations. However we calibrate emergent model dynamics, such as emergent biomasses (used to calibrate $R_{max}$) and emergent response to exploitation (used to calibrate $\varepsilon$). Step by step calibration of the model is described below, and here we describe the procedures for calibrating $\varepsilon$.

Based on the general knowledge of species life-history and expert knowledge, we grouped model species/groups into three categories, reflecting their vulnerability to fishing. We then assumed that these three categories are likely to have following decrease in biomass when low fishing level of 0.2/year (on individuals that are fully recruited to fisheries, assuming knife-edge selectivity at maturation size) was applied to one species at a time (based on rule of thumb expectation in Andersen et al. 2019, Fig. 5.5 and other life-history characteristics, such as longevity). We assumed the following:

Low vulnerability species - *T. caudimaculatus, C. rasor, P. laticlavius, urchins*. When F = 0.2/year, expected decrease in biomass is ca 20-40% (in equilibrium condition).

Medium vulnerability species - *Notolaburs*, *D. lewini*, leatherjackets, *M. freycineti, P. bachus, O. cyanomelas, A. arctidens*, lobsters. When F = 0.2/year expected decrease in biomass is ca 30-50% in equilibrium conditions.

High vulnerability species - *L. forsteri, C. spectabilis, C. laticeps*, boarfish, general predator. When F = 0.2/year expected decrease in biomass is ca 50-75% in equilibrium condition. These species, for example, include a shark *C. laticeps*, long lived site attached *C. spectabilis* and boarfish.

Given these expectations we iteratively tuned $\varepsilon$ and $R_{max}$ values, first fishing each species separately and then fishing all species at once, until species response to 0.2/year fishing mortality was approximately within the desired range and species biomasses were in required relative range (see section 1.4 below). We then calculated emergent fishing level at maximum sustainable yield, by fishing one species at a time with annual fishing mortality ranging from 0.01 to 0.09 (by 0.01), 0.1 to 0.2 (by 0.02), and from 0.25 to 0.95 (by 0.05). Notably, this fishing was applied to one species at a time and does not necessarily reflect resilience to fishing in cases where several species are fishing at once (due to changing nature of species interactions). Fishing selectivity was assumed as knife-edge selectivity at maturation size and does not reflect real fisheries selectivity. Nevertheless, none of the species collapsed, even at very high fishing levels, and all settled into new equilibria biomasses. Emergent response to fishing in some species were higher or lower than general expectations, most likely driven by the dynamic nature of species interactions in the model.

The procedure above produced $F_{msy}$ values indicative of species response to fishing (Table S6), although for low sensitivity species $F_{msy}$ values appeared too high. It is important to note however, that F values applied in *mizer* reflect equilibrium biomass conditions in a size structured model with constant recruitment. These values are therefore quite different from F values estimated in e.g. age-structured stock assessment models, where numbers of individuals in an age class decrease exponentially during a year (but these numbers stay stable in a stable state of a size based model). Therefore F values from *mizer* cannot be directly compared to F values from stock assessments. Because the goal of this study was not to explore fishing responses or species vulnerability, model performance was assumed to be sufficiently good. For future studies, more accurate $F_{msy}$ values should be achieved by further calibrating reproduction allocation functions, maturation size, species interaction matrix, and $\varepsilon$ values, and further analyses should be done to relate size based model F to age structured model F.

**Table S 6.** Calibrating model species reproductive efficiency ε to achieve expected (low, medium, high) vulnerability to fishing.

The emergent $F_{msy}$ values are shown for the main baseline parameter combination (parameter set 1).

| Model species | $\varepsilon$ | $F_{msy}$ | *sensitivity* |
|---|---|---|---|
| Notolabrus | 0.364 | 0.75 | medium |
| L_forsteri | 0.067 | 0.20 | high |
| T_caudimaculatus | 0.418 | 0.95 | low |
| C_rasor | 0.179 | 0.95 | low |
| D_lewini | 0.203 | 0.95 | medium |
| leatherjack | 0.175 | 0.30 | medium |
| C_spectabilis | 0.003 | 0.18 | high |
| C_laticeps | 0.053 | 0.14 | high |
| P_laticlavius | 0.280 | 0.95 | low |
| Boarfish | 0.129 | 0.30 | high |
| M_freycineti | 0.155 | 0.25 | medium |
| P_bachus | 0.114 | 0.50 | medium |
| O_cyanomelas | 0.157 | 0.25 | medium |
| A_arctidens | 0.191 | 0.20 | medium |
| predator | 0.008 | 0.16 | high |
| urchins | 0.116 | 0.95 | low |
| lobsters | 0.210 | 0.30 | medium |

#### 1.6. Model calibration procedure

##### *Initial parameter tuning*

Typically, in multispecies size spectrum (MSS) models optimisation algorithms are used to tune species specific recruitment parameters and resource abundance to observed biomasses or catches (Blanchard et al. 2014). This approach relies on there being a single optimal solution, which is not necessarily the case with complex models. During the model development in this study, we first tried the optimisation approach (with R *optim* function) to find most suitable values for estimated parameters (γ, $\varepsilon$ and $R_{max}$), but algorithms did not converge on a solution and did not produce a system where all species coexisted at expected biomass densities. Therefore, we developed an alternative algorithm to find suitable parameter values. The algorithm does not aim for an optimal solution but rather aims to find a reasonable space by rejecting parameter combinations that produce unrealistic solutions.

This was done in several consecutive steps.

1) Initial parameter values for γ, $\varepsilon$ and $R_{max}$ were set as following.

   $$R_{max,i} = 1000 * \kappa_{mean}\, w_{\infty,i}{}^{-1.5}$$

   following (Blanchard et al. 2014), where $\kappa_{mean}$ is the average resource abundance parameter of the three resource spectra. The multiplier 1000 accounts for the fact that original equation was developed to get recruitment per $m^3$, whereas the model presented here operates at $m^2$ (assuming 500m deep open ocean for which the equation was used), and because coastal areas often have higher productivity than open ocean conditions (additional multiplier of 2).

   The species-specific reproduction efficiency $\varepsilon_i$was set initially as
   $$\varepsilon_i = 0.5\, w_{\infty,i}{}^{-0.5}$$

   which allows for higher reproduction efficiency in small bodied species (Andersen 2019) (Brandl et al. 2019).

   The initial species-specific search rates γ at initial feeding level $f_0$ (of 0.6) where set after Hartvig et al. 2011,

   $$\gamma_i(f_0) = \frac{f_0 h_i \beta_i^{2-\lambda}}{(1-f_0)2\pi\kappa\sigma_i}$$

   but resource λ and $\kappa$ parameters were averaged across three resources. The γ values produced by this equation were too low to sustain growth, possibly because unlike in e.g. North Sea model(Blanchard et al. 2012), only a fraction of background resource is available to fish and because of additional cost of growth assumed in this model. These γ values were therefore multiplied by 3 to ensure sufficient growth and feeding levels. For four predatory species (*D. lewini, P. bachus*, predator, lobsters) these values were multiplied further by 3 to ensure sufficient feeding rates at relatively low prey densities, given that predators did not have much access to the background resource. Note, that assumptions were used to only get initial approximations of parameter values, which were later explored in tuning and uncertainty evaluation (see Fig. S4 for final range of values).

2) Initial species abundances set by default in *mizer* were scaled up or down to achieve relative species biomasses similar to those observed in the system. For this rescaling of initial conditions, the size structure of species was not changed, but only the total abundance.

3) Initial $\varepsilon$ values were then adjusted using the steady() function in *mizer*, which adjusts the reproduction efficiency $\varepsilon$ required to keep model biomasses at the initial condition level, assuming feeding and reproduction level remains at the initial condition state. If the steady() function returned unrealistic $\varepsilon$ values (> 1) we explored other parameters that might affect disproportionate mortality or lack of reproduction in this species, mostly the three non-fixed interaction matrix parameters. While the steady() function bring the system to a steady state at the initial condition level, it does not necessarily lead to system stability and species co-existence through time.

4) To achieve system stability through time $R_{max}$ values for each species were further adjusted in small increments running the model for 150 years iteratively until relative biomasses of all species at the end of the simulation were within the 0.8-1.2 limits of the observed relative biomasses (see "Sequential nudging" section in the code provided in https://github.com/astaaudzi/SEAmodel). If during this tuning stage the model was not approaching a stable state and some species were still going extinct, other parameters had to be revised. This mostly included initial biomasses of species and background spectra (if mortality on background spectra was excessively high), some terms of the interaction matrix (if mortality on some species was extremely high), or intake and metabolism parameters (γ, *h* and $k_s$) if feeding level was sufficiently high (> 0.6) but growth was unrealistically low. Once a desired solution was reached (biomasses were within the set limit, 0.8-1.2 of the observed biomasses in this case), we then calibrated density dependence of recruitment (step 5) and vulnerability to fishing (step 6).

5) We calibrated $\varepsilon$ by aiming to achieve that recruitment in our system is at least partly driven by the spawning stock biomass, and therefore that ratio of realised recruitment *R* (eq 11, TableS1) versus maximum recruitment $R_{max}$ should be within the range of
$R/R_{max}$ ~ [0.85-0.98]
Note, that in some stocks, recruitment might be entirely determined by habitat availability or immigration, and for these cases it could be reasonable to assume that recruitment at equilibrium conditions is very close to the calibrated $R_{max}$ value, but this was not the assumption made in this study.
In addition to the $R/R_{max}$ criterion, we also assessed how a species responds to additional mortality due to fishing. General size-based theory provides expectations on how species with different body sizes are likely to respond to fishing (Fig. S3), or more specifically what is the expected biomass decrease under certain exploitation levels.

6) In the next step, $\varepsilon$ and $R_{max}$ were adjusted iteratively to achieve desired vulnerability to fishing, as described in section 1.5. above.

7) Steps 4, 5 and 6 were repeated several times until both relative biomasses and relative recruitment level are within the desired range. The final recruitment level $R/R_{max}$ was in the range of 0.72-0.99 for various species (can be checked with function getReproductionLevel())

8) Finally, search volume rate (γ) was adjusted further to give daily adult food intake to be between 0.4 to 1.2% of their body weight, as is reported in different estimates (see above). Then steps 4, 6 and 8 were repeated iteratively to arrive at required emergent properties, such as emergent growth rates, feeding level and diets.

#### *Estimating parameter uncertainty using a rejection approach*

The initial steps of parameter uncertainty evaluation are described in the main manuscript (evaluation of 2.2 $x10^6$ parameter combinations against relative equilibrium species biomasses to arrive to a more restricted set of 287 $x10^3$ parameter sets). The second part of parameter rejection followed these steps:

1) Evaluating response to fishing lobsters at a rate of F = 0.2/year, using knife edge selectivity at maturation size. Lobsters are important predators and maintain key control over urchin populations. Reduced lobster abundance from fishing is expected to increase biomasses of some fish compared to the baseline unfished scenarios and also to increase urchin abundance. However this increase or decrease, given the relatively low fishing level, should not be too drastic. We excluded parameter combinations where, after imposing F = 0.2/year fishing on lobsters, relative biomass of at least one species increased or decreased 3-fold (**0.33-3x** range) compared to the unfished baseline scenario. This criterion mostly removed parameter combinations for which lobsters were too sensitive to fishing (decreased too much) or where urchin populations became too abundant (note, the unrealistically strong decrease in lobsters did not always correspond to unrealistically strong increase in urchin populations, as other species also interacted with lobsters and urchins). The criterion reduced the number of parameters *from 287490 to 116108.*

2) Evaluating response to fishing urchins at a rate of F = 0.2/year, using knife edge selectivity at maturation size. Urchins are important ecosystem engineers, although the model here did not include habitat effects on fish, which is the main way how lobsters affect fish (by creating urchins barrens). We therefore do not model species to respond strongly to fishing urchins at a relatively low rate and do not expect large change in the urchin population itself, because urchins are generally resilient to exploitation due to their small size and high productivity. We excluded parameter combinations where relative biomass of at least one species increased or decreased by 50% (**0.67-1.5x** range) compared to the unfished baseline scenario. This criterion mostly filtered cases of unrealistically low or high responses of urchins to fishing, and cases where planktivorous fish showed very strong response to changes in urchin biomass (due to early larval competition for plankton by urchins and planktivores). The criterion reduced the number of parameters *from 116108 to 105720*.

3) Evaluating response to fishing the general predator at a rate of F = 0.2/year, using knife edge selectivity at maturation size. This criterion assessed the importance of top-down control in the ecosystem (including fishing effects on the predator itself). We expect some degree of trophic cascades from reduced predator abundance, but do not expect any species to increase more than two-fold or decrease more than four-fold (**0.25-2x** range). The lower range criterion mostly filters out cases where predators had too strong response to fishing. The criterion reduced the number of parameters *from 105720 to 49815*.

4) Evaluating response to fishing *Trachinops caudimaculatus* (main planktivore fish and important food source for predators) at a rate of F = 0.2/year, using knife edge selectivity at maturation size. While *Trachinops* are not targeted by fishing, they are a key food source and have strong effect on plankton, hence we explore the effect of imposing additional mortality on adults of this species. We expect that the species might respond either positively or negatively to fishing mature individuals (biomass overcompensation can be expected in some species), and but we do not expect the change to be too strong, because small bodied species

are relatively resilient to fishing. We therefore excluded parameter combinations where relative biomass of at least one species increased or decreased by 50% (**0.67-1.5x** range) compared to the unfished baseline scenario. The criterion reduced the number of parameters *from 49815 to 28472* and mostly filtered cases of unrealistically low or high responses of *Trachinops* to fishing, and subsequent effect on other species.

The other three criteria looked at the emergent species characteristics in the unfished system.

5) Feeding level in the unfished system. We assessed feeding level of six keystone species - Notolabrus, predator, Dinolestes, Trachinops, lobsters and urchins. Generally, *mizer* assumes initial feeding level to be at around 0.6 of maximum, satiation level. We therefore only selected parameter combinations where feeding level at maturation size for these groups was within the range of 0.5 and 0.8. The criterion reduced the number of parameters *from 28472 to 1072*. The criterion mostly filtered out cases where planktivore feeding levels were too low.

6) Next, we explored diets of key species at adult body sizes (2/3 of their maximum weight) in the unfished system. Since species diets change through ontogeny and as they grow, we focused on diets of an average adult, for which most information is available. We looked at the proportion of background resources (plankton and benthos) in the diets of key model species (Notolabrus, lobsters, Dinolestes, predators) to ensure that proportions of background resource in predator diets is not too high, as this is often a problem in models and leads to weakened species interactions, because predators end up feeding on resources. Most diets seemed reasonable given the knowledge, however we removed parameter combinations where urchin proportion in the diets of Dinolestes (mostly pelagic predator) exceeded 10% and in the diets of a general predator (big fish, seals, birds) exceeded 20%. The criterion reduced the number of parameters *from 28472 to 255*, filtering out many parameter combinations where proportion of urchins in predator diets was too high.

7) The final criterion applied looked at the biomasses of model species in the unfished scenario and rejected parameter combinations where modelled biomasses ($g/m^2$) were at least two time larger or smaller (**0.5-2x** range) than the observed biomasses over space and time ($g/m^2$). This reduced the parameter set from *255* to the final set of *28* parameter combinations.

The final set of 29 accepted parameter combinations was typically within the 20% range of the initial tuned parameter values.

#### 1.7. Background resource parameters: plankton slope and abundance

##### Plankton slope

For many marine ecosystems the slope of the plankton abundance spectrum varies between -1 and -1.2, which in *mizer* corresponds to -2 and -2.2 ((Edwards et al. 2017, Andersen 2019, Atkinson et al. 2021)). We still lack plankton slope estimates over sufficiently broad size ranges, because slope estimates over fewer than $10^7$ mass ranges are highly variable and less reliable. One of a few long-term empirical plankton slope estimates for a coastal system in the UK (estimated over a large range of mass units) reported slopes that are generally lower than -1.1 (less than -2.1 based on normalised abundance size spectrum in *mizer*), when nitrate and phosphorus concentration were low (Atkinson et al. 2021). Phosphorus and nitrate concentrations for the Maria Island sites, were around 0.3uM and 2.3uM respectively (corresponding to -0.5 and 0.36 on a log10scale), but lower for more northern sites, like Bicheno (-0.57 and 0.28 on a log10 scale) (Stuart-Smith et al. 2013); these values corresponded to the lower end of values reported in (Atkinson et al. 2021), and therefore we used -2.15 for the baseline scenario. Size-based regional (FishMip) models for the off-shore SE Australian systems project that both slope and the total plankton abundance will decrease by 2100 (Fig. S4). While exact values of the slope may differ, the expected change in plankton size spectrum slope is around 0.03, and this is the value we used in our changing slope scenarios. We explore implications of both steeper and shallower slopes, since e.g. recent mesocosm experiment (Barneche et al. 2021) suggest that heated waters will become dominated by larger zooplankton.

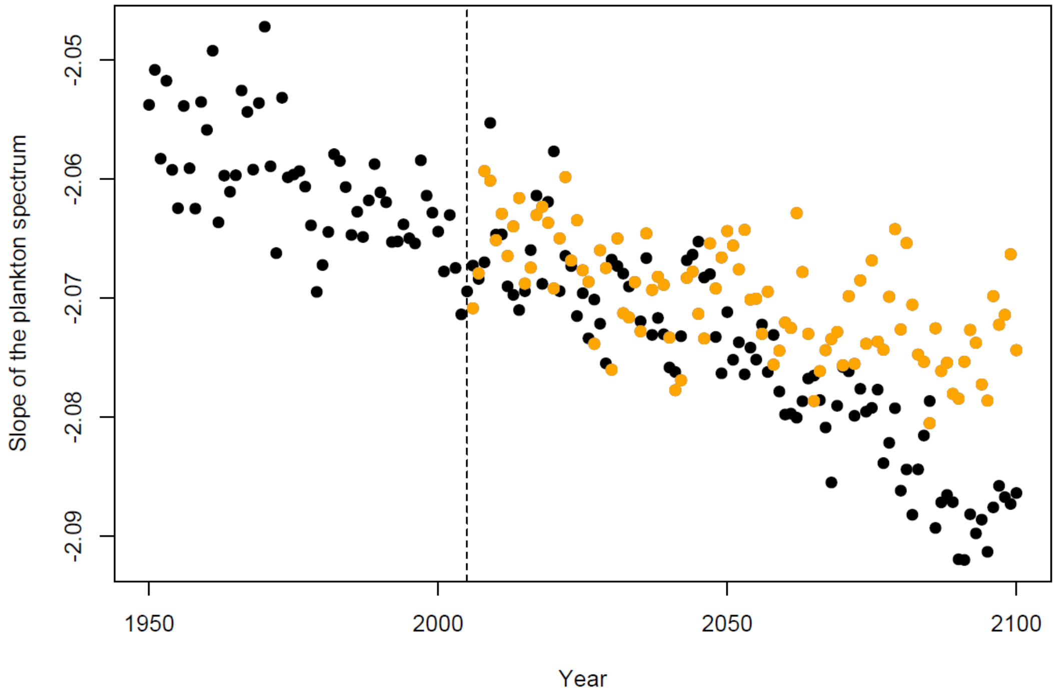

**Figure S 3.** Average annual modelled slopes of plankton size spectrum in off-shore Australian sites for 1950-2100 based on FishMip model projections.

The vertical line separates historical scenarios from future projections, where black dots correspond to RCP85 high emissions scenario used here and orange dots show projections under the RCP45 (lower emissions) scenario.

#### Plankton abundance

The ChlA based estimates of plankton abundance in SE Australian off-shore waters are around 0.03 g/$m^3$ (Bryndum-Buchholz et al. 2019). Assuming the average depth of 100m and given that the model presented here uses $m^2$ and not $m^3$ units, this would translate to ca 3g/$m^2$. The values for coastal reef systems are highly uncertain, because Chla estimates are unreliable in coastal areas. In this study we assume the kappa values of 2 g/$m^2$ in the baseline scenario but explore both higher and lower ranges. When it comes to climate change predictions in coastal plankton abundance, the evidence is even more uncertain. Empirical analyses from 2009-2019 at Maria Island for phytoplankton and zooplankton, did not detect changes in total abundance (and biovolume) for phytoplankton, and a slight increase in abundance (and biomass) for the zooplankton (Everett et al. 2020 Contrasting trends of Australia's plankton communities, State and Trends of Australia's Ocean Report, www.imosoceanreport.org.au). Yet, global model project a nearly 50% decrease in plankton abundance (i.e. intercept in the Fig. S5) by 2100 (Bryndum-Buchholz et al. 2019). We therefore consider both increase and decrease plankton abundance scenarios, allowing the kappa value of the plankton background resource to decrease or increase by 30%.

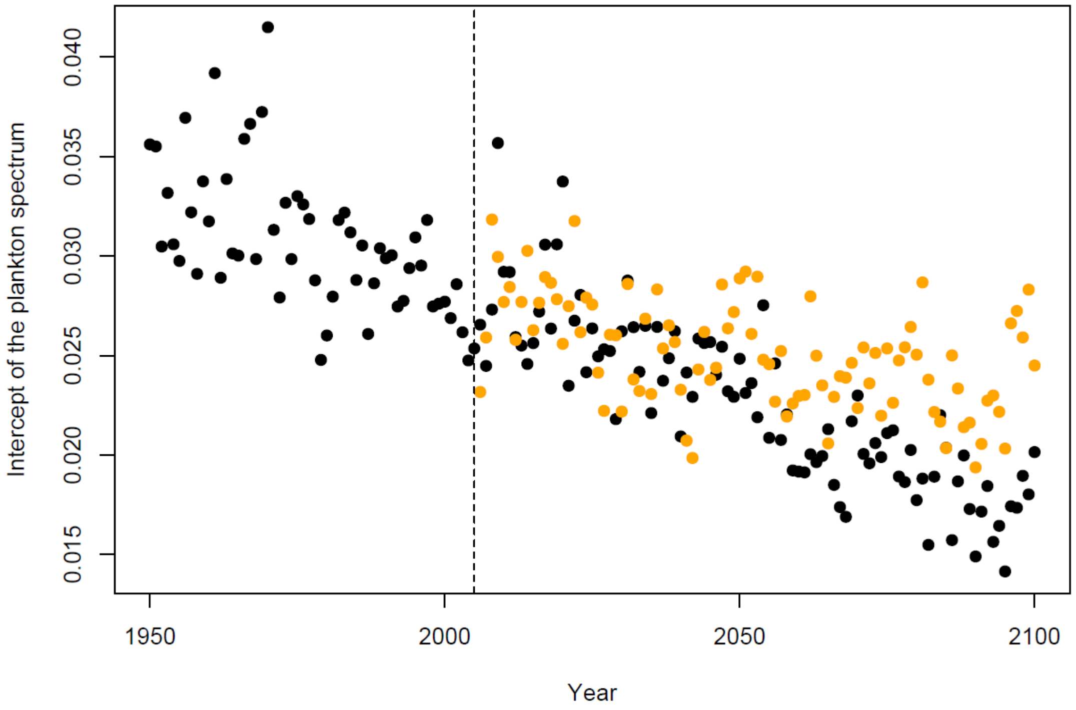

**Figure S 4.** Average annual modelled abundances (intercept) of plankton size spectrum in off-shore Australian sites for 1950-2100 based on FishMip model projections.

##### 1.8. Background resource parameters: benthos slope and abundance

Recent studies of benthic invertebrate abundances along SE Australian coast have been conducted by (Fraser et al. 2021) (using targeted sampling of benthic organisms ranging in size from 1.8e-e06 mg to 15mg) and (Heather et al. 2021a, Heather et al. 2021b) (using underwater visual surveys and invertebrate sizes from 0.3g to 20g). We combined these datasets for SE Tasmanian locations and estimated slope of the benthic spectrum of ca -1.85 for the total data sets, which includes large invertebrates, such as urchins and lobsters (red line in Fig. S6). However, invertebrate over 5g typically represent urchins and lobsters, which are modelled as separate size structured groups in the model presented there. Therefore, for initial analyses we assumed a slightly steeper slope of -1.9 (blue line). The overall abundance of the benthos spectrum (kappa) was iteratively tuned to ensure that the observed abundance across all size groups corresponds to the modelled abundance in the equilibrium conditions (see GitHub code for further details). This gave the kappa value of 6 $g/m^2$. Benthos abundance in coastal systems generally is expected to be more variable and responsive than plankton abundance (which is more affected by the influx from offshore systems), hence we allowed more variation in response to climate change and allowed slope changes of 0.1 and kappa increase up to 9 $g/m^2$ or decrease to 4 $g/m^2$ (ca 30-50%).

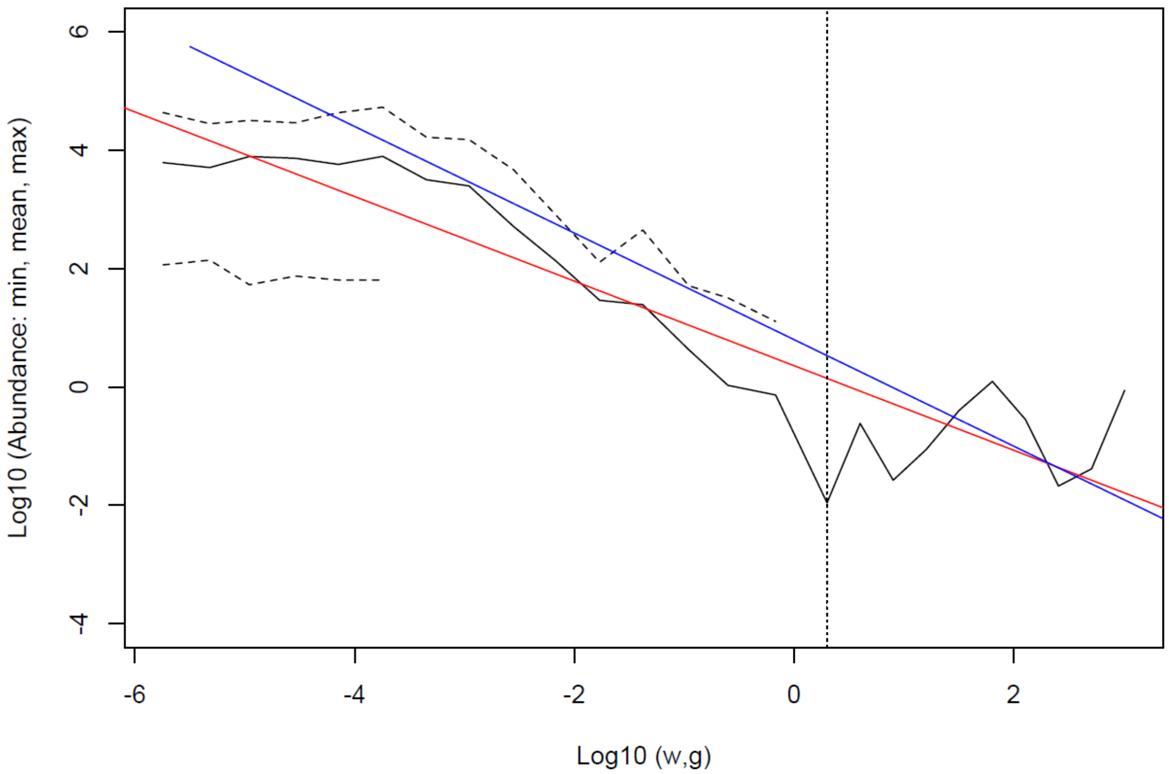

**Figure S 5.** Observed (black line and dashed lines) abundances of benthic invertebrates in eastern Tasmania

Data from (Fraser et al. 2021) and (Heather et al. 2021a, Heather et al. 2021b) shown with black lines. We explored a range of slopes values and assessed their fit to data visually. A slope of -1.85 is shown with a red line, slope of -1.9 with a blue line. See GitHub code for data and further details.

#### 2. Results

##### 2.1. Baseline biomass variation

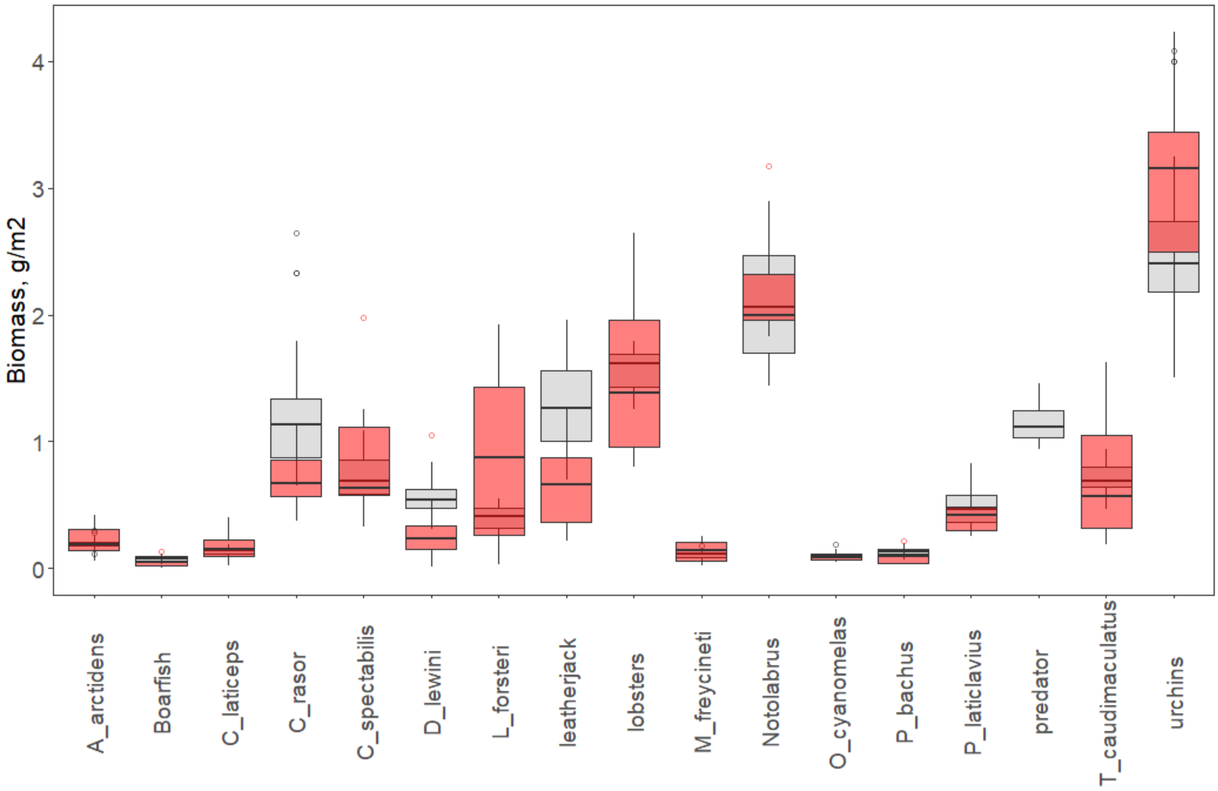

**Figure S 6.** Modelled and observed biomasses.

Modelled (grey boxes) variation in equilibrium biomasses (g/m$^2$) of the 17 model species and groups in 29 parameter combinations, in baseline scenario of no plankton and benthos resource change and low fishing mortality (Table 1). Red boxes show interannual variation in the observed mean annual biomasses in visual surveys during 1992-1999. Biomasses of urchins and lobsters were estimated based on visual survey abundance estimates assuming an average lobster weight of 150g and an average urchin weight of 5g. Predator is an artificial group combining all predation not specifically observed in visual surveys.

##### 2.2. Changes in the resource slopes and impacts on equilibrium biomasses

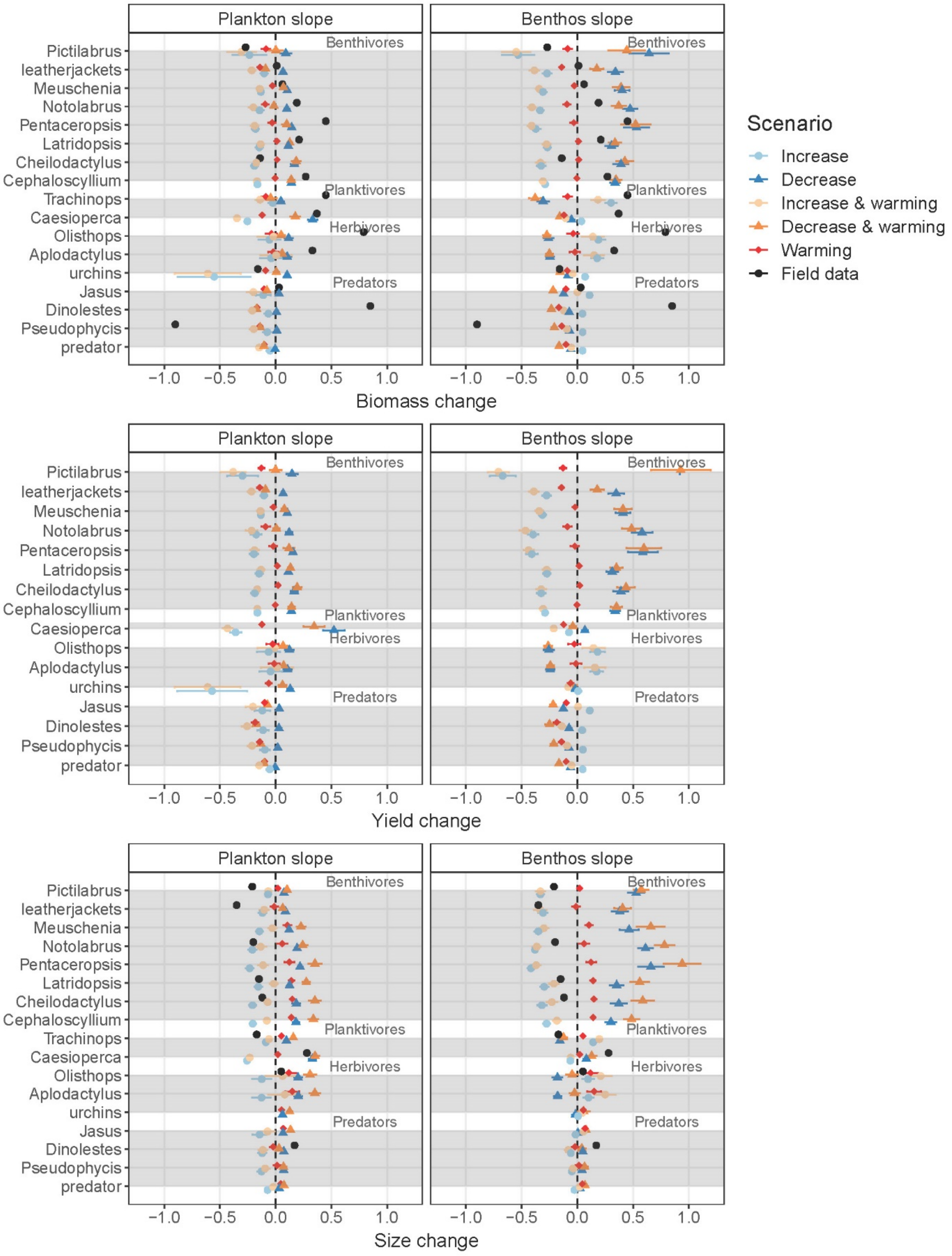

**Figure S 7.** Changes in biomasses, yields and mean size in response to changing plankton and benthos size spectrum slopes. Changes in biomasses, yields and mean body sizes (individuals above 2cm length) in model groups across the alternative scenarios of steeper or shallower plankton and benthos resource slopes. Variation across all 29 parameter combinations is shown with error bars. For other details see explanations in Figure 3.

#### 2.3. Mixed effect model selection and results

For most analyses the best model included all two-way interactions, but not the three-way interaction. The only exception was plankton change effect on species mean size, where one two-way interaction between warming and functional group was not significant.

**Table S 7.** Selection of models for mixed-effect ANOVA analyses.

Here we used R function drop1() to explore alternative scenarios. “Bck” refers to background resource, “fnc” is functional group, “wrm” is warming. In most cases, except for benthos and plankton effect on mean body size, the best scenario had two- or three-way interactions between all three parameters (marked in red). For consistency we used two-way interaction models in further analyses.

##### Benthos kappa changes and biomass

```
Global model call: lmer(formula = value ~ 1 + funcgr * backgr * warming + (1 | species),
    data = df_be_bio, REML = T)
---
Model selection table
     (Int) bck fnc wrm bck:fnc bck:wrm fnc:wrm bck:fnc:wrm df   logLik   AICc   delta weight
64   1.2270  +   +   +       +       +       +             20 1931.287 -3822.3   0.00  0.968
48   1.2400  +   +   +       +               +             18 1925.855 -3815.5   6.81  0.032
128  1.2220  +   +   +       +       +       +           + 26 1926.617 -3800.8  21.53  0.000
32   1.2420  +   +   +       +       +                     17 1912.279 -3790.3  31.94  0.000
16   1.2550  +   +   +       +                             15 1907.066 -3784.0  38.32  0.000
```

##### Benthos kappa changes and yields

```
Global model call: lmer(formula = value ~ 1 + funcgr * backgr * warming + (1 | species),
    data = df_be_yield, REML = F)
---
Model selection table
     (Int) bck fnc wrm bck:fnc bck:wrm fnc:wrm bck:fnc:wrm df    logLik   AICc   delta weight
64   1.2300  +   +   +       +       +       +             20   362.995 -685.7   0.00  0.968
128  1.2270  +   +   +       +       +       +           + 26   365.673 -678.8   6.85  0.031
48   1.2500  +   +   +       +               +             18   353.405 -670.6  15.12  0.001
32   1.2530  +   +   +       +       +                     17   350.701 -667.2  18.51  0.000
```

##### Benthos kappa changes and biomass

```
Global model call: lmer(formula = value ~ 1 + funcgr * backgr * warming + (1 | species),
    data = df_be_size, REML = F)
---
Model selection table
     (Int) bck fnc wrm bck:fnc bck:wrm fnc:wrm bck:fnc:wrm df   logLik   AICc   delta weight
128 1.1680  +   +   +       +       +       +           + 26 2579.922 -5107.4   0.00      1
48   1.2040  +   +   +       +               +             18 2558.262 -5080.3  27.07      0
64   1.1990  +   +   +       +       +       +             20 2560.214 -5080.1  27.22      0
16   1.2240  +   +   +       +                             15 2517.592 -5005.0 102.34      0
```

##### Plankton kappa changes and biomass

```
Global model call: lmer(formula = value ~ 1 + funcgr * backgr * warming + (1 | species),
    data = df_pl_bio, REML = T)
---
Model selection table
     (Int) bck fnc wrm bck:fnc bck:wrm fnc:wrm bck:fnc:wrm df   logLik   AICc   delta weight
64   0.7639  +   +   +       +       +       +             20 1791.089 -3541.9   0.00  0.785
48   0.7786  +   +   +       +               +             18 1787.770 -3539.3   2.59  0.215
128  0.7419  +   +   +       +       +       +           + 26 1780.599 -3508.7  33.17  0.000
32   0.7832  +   +   +       +       +                     17 1765.959 -3497.7  44.18  0.000
```

##### Plankton kappa changes and yields

```
Global model call: lmer(formula = value ~ 1 + funcgr * backgr * warming + (1 | species),
    data = df_pl_yield, REML = T)
---
Model selection table
     (Int) bck fnc wrm bck:fnc bck:wrm fnc:wrm bck:fnc:wrm df   logLik   AICc   delta weight
64   0.7206  +   +   +       +       +       +             20 2001.611 -3962.9   0.00      1
128  0.7009  +   +   +       +       +       +           + 26 1996.343 -3940.2  22.74      0
48   0.7423  +   +   +       +               +             18 1981.876 -3927.5  35.41      0
32   0.7375  +   +   +       +       +                     17 1967.275 -3900.3  62.59      0
```

##### Plankton kappa changes and mean body size

```
Global model call: lmer(formula = value ~ 1 + funcgr * backgr * warming + (1 | species),
    data = df_pl_size, REML = T)
---

     (Int) bck fnc wrm bck:fnc bck:wrm fnc:wrm bck:fnc:wrm df    logLik    AICc  delta weight
12  13.970   +   +           +                             14 -9793.080 19614.3   0.00  0.656
16  13.970   +   +   +       +                             15 -9793.578 19617.3   3.02  0.145
128 14.510   +   +   +       +       +       +           + 26 -9782.708 19617.9   3.59  0.109
32  14.030   +   +   +       +       +                     17 -9792.850 19619.9   5.60  0.040
64  14.180   +   +   +       +       +       +             20 -9791.136 19622.6   8.25  0.011
4    6.745   +   +                                          8 -9976.136 19968.3 354.02  0.000
```

**Table S 8.** Coefficients from the final mixed effect ANOVA analysis, with their 95% confidence intervals.

**Benthos change**

| BIOMASS | Estimate | 2.5%CI | 97.5%CI | t-value |
|---|---|---|---|---|
| (Intercept) | 1.227 | 1.194 | 1.26 | 67.297 |
| benthivore | -0.586 | -0.625 | -0.547 | -27.839 |
| planktivore | -0.034 | -0.086 | 0.018 | -1.207 |
| predator | -0.325 | -0.368 | -0.281 | -13.683 |
| ResourceStable | -0.231 | -0.259 | -0.203 | -16.148 |
| ResourceIncrease | -0.399 | -0.427 | -0.371 | -27.93 |
| warming | -0.018 | -0.043 | 0.006 | -1.442 |
| benthivore:Resource Stable | 0.592 | 0.561 | 0.622 | 38.297 |
| planktivore:Resource Stable | 0.037 | -0.003 | 0.078 | 1.797 |
| predator:Resource Stable | 0.328 | 0.294 | 0.362 | 18.818 |
| benthivore:Resource Increased | 1.21 | 1.179 | 1.24 | 78.304 |
| planktivore:Resource Increased | 0.017 | -0.024 | 0.058 | 0.817 |
| predator:Resource Increased | 0.611 | 0.577 | 0.645 | 35.058 |
| benthivore:warming | -0.007 | -0.031 | 0.018 | -0.522 |
| planktivore:warming | -0.061 | -0.094 | -0.028 | -3.598 |
| predator:warming | -0.084 | -0.112 | -0.056 | -5.889 |
| ResourceStable:warming | -0.025 | -0.046 | -0.003 | -2.214 |
| ResourceIncrease:warming | -0.056 | -0.077 | -0.034 | -5.046 |

| YIELD | Estimate | 2.5%CI | 97.5%CI | t-value |
|---|---|---|---|---|
| (Intercept) | 1.23 | 1.163 | 1.297 | 37.199 |
| benthivore | -0.624 | -0.701 | -0.546 | -16.333 |
| planktivore | -0.061 | -0.193 | 0.072 | -0.932 |
| predator | -0.36 | -0.447 | -0.272 | -8.349 |
| ResourceStable | -0.235 | -0.284 | -0.187 | -9.547 |
| ResourceIncrease | -0.401 | -0.449 | -0.353 | -16.267 |
| warming | 0.013 | -0.03 | 0.055 | 0.586 |
| benthivore:Resource Stable | 0.633 | 0.581 | 0.685 | 23.844 |
| planktivore:Resource Stable | 0.066 | -0.023 | 0.154 | 1.453 |
| predator:Resource Stable | 0.363 | 0.305 | 0.422 | 12.142 |
| benthivore:Resource Increased | 1.401 | 1.349 | 1.453 | 52.816 |
| planktivore:Resource Increased | 0.094 | 0.005 | 0.183 | 2.075 |

| | | | | |
|---|---|---|---|---|
| predator:Resource Increased | 0.688 | 0.629 | 0.747 | 22.984 |
| benthivore:warming | -0.03 | -0.072 | 0.013 | -1.37 |
| planktivore:warming | -0.1 | -0.173 | -0.028 | -2.709 |
| predator:warming | -0.105 | -0.153 | -0.057 | -4.296 |
| ResourceStable:warming | -0.035 | -0.074 | 0.003 | -1.81 |
| ResourceIncrease:warming | -0.086 | -0.124 | -0.047 | -4.366 |

| **MEAN BODY SIZE** | Estimate | 2.5%CI | 97.5%CI | t-value |
|---|---|---|---|---|
| (Intercept) | 1.199 | 1.162 | 1.237 | 65.332 |
| benthivore | -0.523 | -0.567 | -0.48 | -24.564 |
| planktivore | -0.096 | -0.155 | -0.038 | -3.358 |
| predator | -0.35 | -0.399 | -0.301 | -14.559 |
| ResourceStable | -0.202 | -0.225 | -0.179 | -17.189 |
| ResourceIncrease | -0.353 | -0.376 | -0.33 | -30.032 |
| warming | 0.124 | 0.104 | 0.144 | 12.031 |
| benthivore:Resource Stable | 0.526 | 0.502 | 0.551 | 41.438 |
| planktivore:Resource Stable | 0.101 | 0.067 | 0.134 | 5.877 |
| predator:Resource Stable | 0.353 | 0.325 | 0.381 | 24.646 |
| benthivore:Resource Increased | 1.135 | 1.11 | 1.16 | 89.307 |
| planktivore:Resource Increased | 0.153 | 0.12 | 0.187 | 8.945 |
| predator:Resource Increased | 0.697 | 0.669 | 0.725 | 48.62 |
| benthivore:warming | -0.023 | -0.043 | -0.003 | -2.223 |
| planktivore:warming | -0.08 | -0.108 | -0.053 | -5.752 |
| predator:warming | -0.088 | -0.11 | -0.065 | -7.479 |
| ResourceStable:warming | -0.011 | -0.029 | 0.007 | -1.177 |
| ResourceIncrease:warming | -0.018 | -0.036 | 0 | -1.964 |

### Plankton change

| **BIOMASS** | Estimate | 2.5%CI | 97.5%CI | t-value |
|---|---|---|---|---|
| (Intercept) | 0.764 | 0.686 | 0.842 | 17.841 |
| benthivore | 0.047 | -0.044 | 0.137 | 0.929 |
| planktivore | -0.067 | -0.19 | 0.056 | -0.991 |
| predator | 0.096 | -0.006 | 0.199 | 1.703 |
| ResourceStable | 0.228 | 0.199 | 0.257 | 15.281 |
| ResourceIncrease | 0.294 | 0.265 | 0.323 | 19.693 |
| warming | 0.003 | -0.023 | 0.028 | 0.201 |
| benthivore:Resource Stable | -0.038 | -0.069 | -0.006 | -2.335 |
| planktivore:Resource Stable | 0.082 | 0.04 | 0.125 | 3.781 |
| predator:Resource Stable | -0.087 | -0.123 | -0.052 | -4.793 |
| benthivore:Resource Increased | 0.01 | -0.021 | 0.042 | 0.63 |
| planktivore:Resource Increased | 0.38 | 0.337 | 0.422 | 17.447 |
| predator:Resource Increased | -0.12 | -0.156 | -0.085 | -6.603 |
| benthivore:warming | -0.013 | -0.039 | 0.013 | -0.991 |
| planktivore:warming | -0.086 | -0.12 | -0.051 | -4.817 |
| predator:warming | -0.096 | -0.125 | -0.067 | -6.435 |

| | | | | |
|---|---|---|---|---|
| ResourceStable:warming | -0.036 | -0.059 | -0.014 | -3.155 |
| ResourceIncrease:warming | -0.052 | -0.074 | -0.029 | -4.476 |

| **YIELD** | Estimate | 2.5%CI | 97.5%CI | t-value |
|---|---|---|---|---|
| (Intercept) | 0.751 | 0.671 | 0.831 | 16.978 |
| benthivore | 0.034 | -0.06 | 0.127 | 0.654 |
| planktivore | -0.418 | -0.578 | -0.259 | -4.742 |
| predator | 0.078 | -0.028 | 0.183 | 1.332 |
| ResourceStable | 0.243 | 0.215 | 0.27 | 17.474 |
| ResourceIncrease | 0.32 | 0.293 | 0.347 | 23.048 |
| warming | 0.029 | 0.005 | 0.053 | 2.357 |
| benthivore:Resource Stable | -0.026 | -0.055 | 0.003 | -1.759 |
| planktivore:Resource Stable | 0.437 | 0.387 | 0.487 | 17.15 |
| predator:Resource Stable | -0.07 | -0.103 | -0.037 | -4.175 |
| benthivore:Resource Increased | 0.033 | 0.004 | 0.062 | 2.228 |
| planktivore:Resource Increased | 1.56 | 1.51 | 1.61 | 61.253 |
| predator:Resource Increased | -0.098 | -0.131 | -0.065 | -5.823 |
| benthivore:warming | -0.027 | -0.051 | -0.003 | -2.23 |
| planktivore:warming | -0.127 | -0.168 | -0.087 | -6.118 |
| predator:warming | -0.112 | -0.139 | -0.086 | -8.172 |
| ResourceStable:warming | -0.049 | -0.071 | -0.028 | -4.461 |
| ResourceIncrease:warming | -0.072 | -0.094 | -0.051 | -6.558 |

| **MEAN BODY SIZE** | Estimate | 2.5%CI | 97.5%CI | t-value |
|---|---|---|---|---|
| (Intercept) | 0.932 | 0.897 | 0.967 | 48.907 |
| benthivore | -0.164 | -0.205 | -0.124 | -7.47 |
| planktivore | -0.237 | -0.291 | -0.183 | -8.044 |
| predator | -0.123 | -0.168 | -0.078 | -4.978 |
| ResourceStable | 0.054 | 0.031 | 0.076 | 4.701 |
| ResourceIncrease | 0.2 | 0.178 | 0.222 | 17.518 |
| warming | 0.166 | 0.146 | 0.185 | 16.77 |
| benthivore:Resource Stable | 0.181 | 0.157 | 0.205 | 14.791 |
| planktivore:Resource Stable | 0.256 | 0.225 | 0.288 | 15.856 |
| predator:Resource Stable | 0.141 | 0.114 | 0.167 | 10.279 |
| benthivore:Resource Increased | 0.222 | 0.199 | 0.246 | 18.18 |
| planktivore:Resource Increased | 0.521 | 0.49 | 0.553 | 32.249 |
| predator:Resource Increased | 0.094 | 0.068 | 0.121 | 6.905 |
| benthivore:warming | -0.05 | -0.069 | -0.03 | -5.078 |
| planktivore:warming | -0.111 | -0.136 | -0.085 | -8.476 |
| predator:warming | -0.115 | -0.137 | -0.094 | -10.522 |
| ResourceStable:warming | -0.03 | -0.047 | -0.014 | -3.561 |
| ResourceIncrease:warming | -0.035 | -0.051 | -0.018 | -4.141 |
